# Structural and Mechanistic Basis of F227C-Mediated Hypersusceptibility to Islatravir in HIV-1 Reverse Transcriptase

**DOI:** 10.64898/2026.06.08.730881

**Authors:** Alexa A. Snyder, Isabella L. Kaufman, Jack MacQuillan, Ryan L. Slack, Karen A. Kirby, Eleftherios Michailidis, Stefan G. Sarafianos

**Affiliations:** Center for ViroScience and Cure, Laboratory of Biochemical Pharmacology, Department of Pediatrics, Emory University School of Medicine, Atlanta, GA 30322; Children’s Healthcare of Atlanta, Atlanta, GA 30322

## Abstract

Islatravir (ISL; 4′-ethynyl-2-fluoro-2′-deoxyadenosine (EFdA)), a first-in-class nucleoside reverse transcriptase (RT) translocation inhibitor (NRTTI), was recently approved by the FDA for the treatment of HIV-1 infection in combination with the non-nucleoside RT inhibitor (NNRTI), doravirine (DOR). Notably, the RT mutation F227C, which confers clinical resistance to multiple NNRTIs, including DOR, unexpectedly increases susceptibility to ISL. To elucidate the mechanistic basis of this hypersusceptibility, we determined a 1.8 angstrom crystal structure of F227C RT in complex with a ddGMP-terminated primer/template and ISL-triphosphate. The structure reveals conformational rearrangements that propagate into a cleft, thereby affecting ATP-mediated unblocking of chain-terminating antivirals. Complementary biochemical assays showed that although F227C does not significantly affect ISL incorporation, it alters RT translocation and impairs ATP-dependent phosphorolytic excision of ISL-terminated primers, thereby enhancing ISL susceptibility. These findings establish direct structural and mechanistic links between NNRTI resistance and ISL hypersusceptibility, providing a structural foundation for rationally designed, resistance-informed combination regimens that exploit this unique collateral sensitivity.

## Introduction

As of 2024, approximately 40.8 million people are living with human immunodeficiency virus (HIV), with 72% of these people achieving viral suppression through antiretroviral therapy (ART) (1). Despite this success, 1.3 million new infections and over 630,000 Acquired Immunodeficiency Syndrome (AIDS)-related deaths occurred in the past year (1). Limitations in current regimens, driven by adherence challenges, drug resistance, and access barriers, continue to fuel the development of next-generation antiretrovirals, including long-acting agents, novel mechanisms of action, and optimized combinations (2-5).

HIV therapies almost invariably combine multiple drugs that target distinct steps in the viral replication. Reverse transcriptase (RT) remains a fundamental target because it is essential for converting the viral single-stranded RNA genome (+ssRNA) into double-stranded DNA (dsDNA) that is subsequently integrated into the host genome (6). Three major classes of approved antivirals target RT: nucleoside reverse transcriptase inhibitors (NRTIs), nucleoside reverse transcriptase translocation inhibitors (NRTTI), and non-nucleoside reverse transcriptase inhibitors (NNRTIs). NRTIs and NRTTIs are chain-terminating nucleoside analogs that compete with natural dNTPs at the polymerase active site in the p66 subdomain. Classic NRTIs lack a 3’-OH and block RT through obligate termination once incorporated into the elongating DNA. In contrast, NRTTIs retain the 3’-OH and can inhibit through immediate or delayed chain termination (7). NNRTIs bind a distinct hydrophobic pocket located at the base of the p66 thumb subdomain (8, 9), which is only formed in the presence of an NNRTI (10, 11). NNRTIs restrict the mobility of the p66 thumb, altering the positioning of the primer and/or restricting the flexibility of the Y183-M184-D185-D186 (YMDD) loop that is a critical part of the catalytic site of RT, leading to inhibition of DNA synthesis (7, 12-15). Despite the success of RT inhibitors, the efficacy of these antivirals can be significantly reduced by drug-resistant mutations.

Islatravir, (ISL; 4′-ethynyl-2-fluoro-2′-deoxyadenosine (EFdA)) is a first-in-class NRTTI recently approved in combination with doravirine (DOR), as Idvynso (16). This two-drug combination is now the first and only non-INSTI (Integrase Strand Transfer Inhibitor), tenofovir-free therapy for the treatment of HIV infection, shown to be non-inferior to the most potent three-drug regimens, including Biktarvy (16, 17). Unlike conventional NRTIs, ISL inhibits DNA synthesis by the immediate or delayed blocking of RT translocation and increased misincorporation and it was thus termed an NRTTI. Its 4′-ethynyl and 2-fluoro modifications, combined with the retained 3′-OH, confer exceptional potency (EC_50_ ∼0.07 nM), a long intracellular half-life (78.5-128 hours), and a high selectivity index (18-21). These properties support its potential for long-acting regimens.

Multiple NRTI resistance mutations (RMs) have been reported at RT residues 41, 65, 67, 68, 69, 70, 74, 115, 151, 184, 210, and 215 of RT (7, 22-24). These RMs confer resistance primarily through NRTI discrimination or enhanced excision (6, 7, 25, 26). Common discrimination mutations include M184V/I, primarily affecting 3TC and FTC; K65R, affecting tenofovir (TDF), 3TC, FTC, and (ABC), K70E, affecting ABC and TDF, L74V/I primarily affecting ABC, and Y115F, affecting ABC and TDF susceptibility (27). RMs M41L, D67N, K70R, T215F/Y, and K291Q/E increase RT’s excision function, which is an ATP-mediated phosphorolytic activity that unblocks inhibitor-terminated primers (25, 27-31). Passaging experiments identified the first mutations that cause ISL resistance (18). Combined, these mutations result in moderate ISL resistance (22-fold increase in EC_50_). Individually, they cause low-level or no resistance: M184V (7.5-fold), I142V (0.9-fold), and T165R (1.5-fold resistance). The triple mutant virus has impaired replication fitness and has not been seen in the clinic. Another study showed that rilpivirine (RPV) RM E138K is a compensatory mutation for ISL resistance, as its pairing with M184V/I increased viral fitness compared to M184V/I (32). It was also shown that M41L/M184V/A114S and M184V/A114S, and M184V/V106I reduce the ISL efficacy by 24-to 65-fold, but they also imparted low viral fitness (33-35). Overall, ISL maintains a high genetic barrier to resistance.

Some RMs cause hypersusceptibility to other agents, such as the TDF RM K65R, which we have shown to increase susceptibility to ISL (36). The NNRTI RM F227C, which is not located in the catalytic active site, has been reported to confer increased susceptibility to ISL (37). F227C confers substantial resistance to DOR, an FDA-approved NNRTI, and was identified as one of the most common RMs observed in the DRIVE clinical trials (NCT02275780, NCT02403674, NCT02629822, and NCT02397096) (38-41), frequently occurring alongside V106A/I/M. Residue F227 is located in the p66 palm subdomain of RT, near the p66 thumb hinge region. DOR binds within the non-nucleoside inhibitor-binding pocket (NNIBP) and interacts with K103, V106, Y188, P225, F227, and W229 (PDB: 4NCG) (42). The structural mechanism by which F227C enhances ISL susceptibility, despite being >16 Å from the ISL-triphosphate (TP) binding site, remains unknown. Recent kinetic analysis of the incorporation efficiency of ISL-TP and dATP suggested that F227C RT incorporates the natural substrate dATP 0.6-fold less efficiently than ISL-TP, whereas WT RT incorporates dATP 1.3-fold the efficiency of ISL-TP (43). However, these differences cannot fully account for the reduction in EC_50_ relative to the WT virus (11.9-fold in Figure 1). Thus, the structural and detailed mechanistic basis of F227C RT to ISL remains unclear.

**Figure 1.**
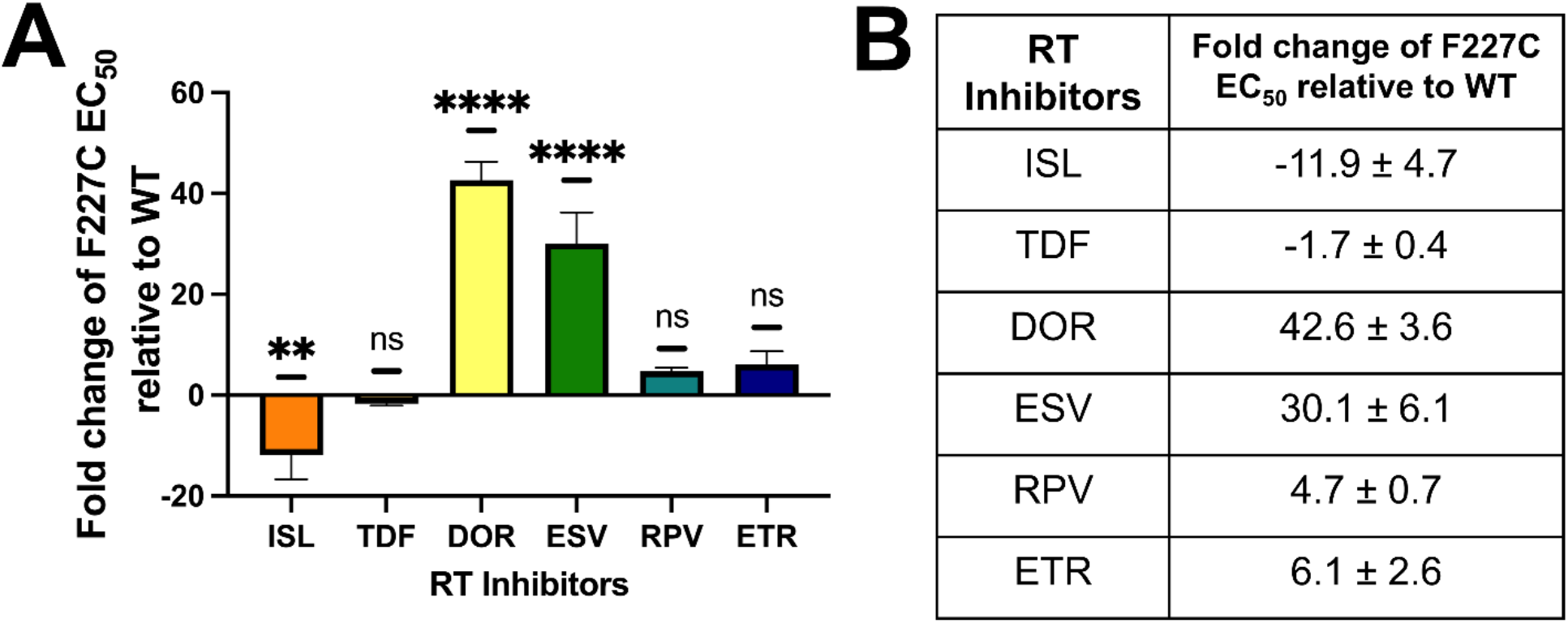
Resistance and hypersusceptibility of HIV-1 F227C pNL4-3Δenv VSV-G pseudotyped virus compared to HIV-1 WT pNL4-3Δenv VSV-G pseudotyped virus against a panel of RT inhibitors. (A) Dose-response experiments were conducted by infecting TZM-GFP cells with pNL4-3Δenv VSV-G pseudotyped WT and F227C mutant virus in the presence of RT inhibitors at various concentrations. The EC_50_ values were calculated in GraphPad Prism using a four-parameter variable-slope nonlinear regression model [log(inhibitor) vs. normalized response] and normalized to WT. Values below 1 were considered hypersusceptible and converted to -1/fold-change for representation. Statistical significance was determined using one-way ANOVA with multiple comparisons (**** P<0.0001, ** P<0.01, ns = not significant). (B) The mean and standard deviation (SD) of the fold change in F227C NL4-3Δenv VSV-G EC_50_ values relative to WT NL4-3Δenv VSV-G virus (n=3-5).

Here, we determined the 1.8 Å crystal structure of the F227C RT in complex with a ddGMP-terminated primer/template and ISL-triphosphate (TP) (RT_F227C_/DNA_ddGMP_/ISL-TP). Combined with biochemical assays, steady-state kinetic analyses of nucleotide incorporation, ATP-mediated phosphorolytic excision of ISL-terminated primers, and hydroxyl radical footprinting experiments, we show that the preliminary, phenotype-causing effect of F227C is impaired RT excision. We also evaluated the therapeutic potential of combining ISL with NNRTIs affected by F227C-mediated resistance. These findings reveal a direct mechanistic link between NNRTI resistance and NRTTI hypersusceptibility, providing a structural framework for exploiting this collateral sensitivity in resistance-informed combination therapies.

## Results

### F227C confers hypersusceptibility to ISL while producing high-level resistance to multiple NNRTIs

To define the antiviral profile of the F227C mutant, we generated HIV-1 pNL4-3 ΔEnv VSV-G pseudotyped viruses and measured susceptibility to a panel of clinically relevant RT inhibitors in TZM-GFP reporter cells (Figure 1). The F227C variant displayed pronounced hypersusceptibility to ISL, with an 11.9-fold reduction in EC_50_ relative to wild-type (WT) virus (Figure 1A, B, Supplemental Figure 1), consistent with previous reports (37, 44). This effect was selective: susceptibility to the adenosine analog tenofovir disoproxil fumarate (TDF) was only modestly increased (1.7-fold decrease in EC_50_). F227C conferred high-level resistance to NNRTIs DOR (42.6-fold) and elsulfavirine (ESV; 30.1-fold), with more modest, non-statistically significant reductions in potency against rilpivirine (RPV) and etravirine (ETR) (Figure 1A, B, Supplemental Figure 1). These data establish that F227C simultaneously confers robust resistance to multiple NNRTIs while creating a striking collateral hypersusceptibility to ISL.

### F227C impairs ATP-dependent rescue of ISL-terminated primers without substantially altering incorporation

We next used recombinant enzymes RT_WT_ and RT_F227C_ to dissect the biochemical basis of ISL hypersusceptibility. Under standard primer-extension conditions lacking ATP, the two enzymes showed nearly identical sensitivity to ISL-TP (non-statistically significant 1.1-fold change in IC_50_ [Half-Maximal Inhibitory Concentration]) (Figure 2A, B). However, in gel-based assays, the addition of physiological concentrations of ATP (Figure 2A, B, C) revealed hypersusceptibility in the mutant, with an IC_50_ fold change of 0.6. These results provided initial evidence that ATP-dependent rescue mechanisms are less efficient in the F227C background. Primer extension experiments were further analyzed to understand changes in inhibition patterns by ISL-TP (Figure 2C). ISL, an NRTTI, disrupts RT translocation in a sequence-dependent manner, acting as either as an obligate chain terminator or as a delayed chain terminator. Consistent with previous studies, strong inhibition was observed at the P+1 position, the classical obligate (45). No major differences between WT and F227C RT were observed in this position (Figure 3C). In contrast, differences emerged at the downstream P+6 “slippery site,” where WT RT more efficiently extended beyond the ISL-MP terminus, whereas F227C RT showed increased accumulation of the P+6 product and reduced progression to P+7, indicating decreased translocation efficiency. This translocation defect was further enhanced in the presence of ATP (Figure 2C).

**Figure 2.**
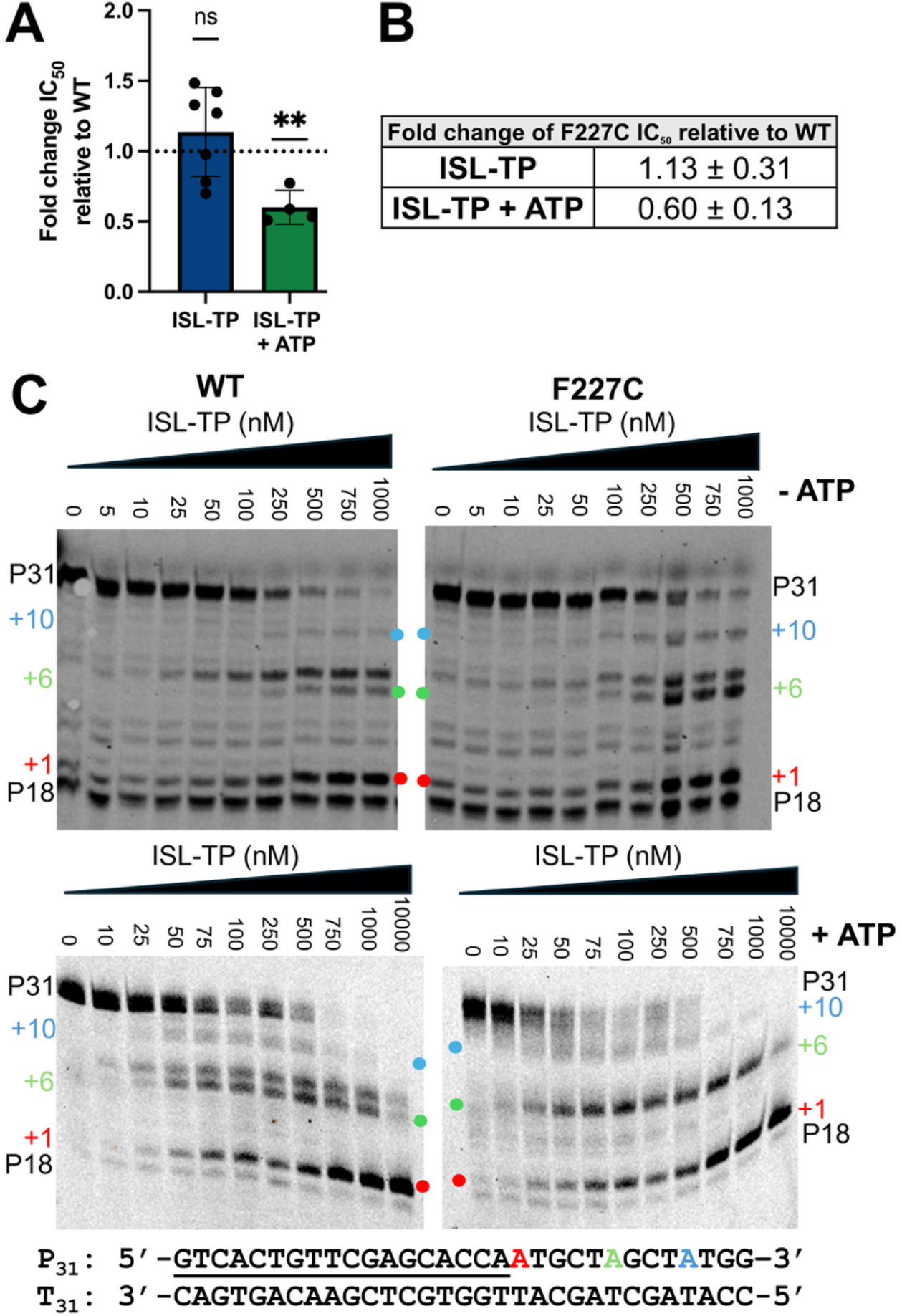
ISL resistance and hypersusceptibility of HIV-1 RT_F227C_ compared with RT_WT_ in the presence or absence of ATP. The Td_31_/Pd_18_-P_0_ oligonucleotide substrate was incubated with RT_WT_ or RT_F227C_ in the presence of 1 μM dNTPs, 6 mM MgCl_2_, and increasing concentrations of islatravir triphosphate (ISL-TP) (0–10,000 nM), with or without 3.5 mM ATP. The 3′ end of the annealed Pd_18_ primer (underlined above) corresponds to the P_0_ site. (A) Percent primer extension was plotted across the ISL-TP concentration range, and IC_50_ values were calculated in GraphPad Prism using a four-parameter variable-slope nonlinear regression model [log(inhibitor) vs. normalized response] and normalized to RT_WT_. (B) Normalized IC_50_ values were plotted and analyzed using a one-sample t-test and Wilcoxon signed-rank test (*P < 0.05). (C) Reaction products were resolved on a 15% polyacrylamide/7 M urea gel. The template sequence is shown below the gel, and numbers indicate ISL-TP incorporation and translocation sites (+1, +6, and +10). Colored dots indicate the corresponding positions at the opposite end of the gel (n = 7 for −ATP assays; n = 4 for +ATP assays).

**Figure 3.**
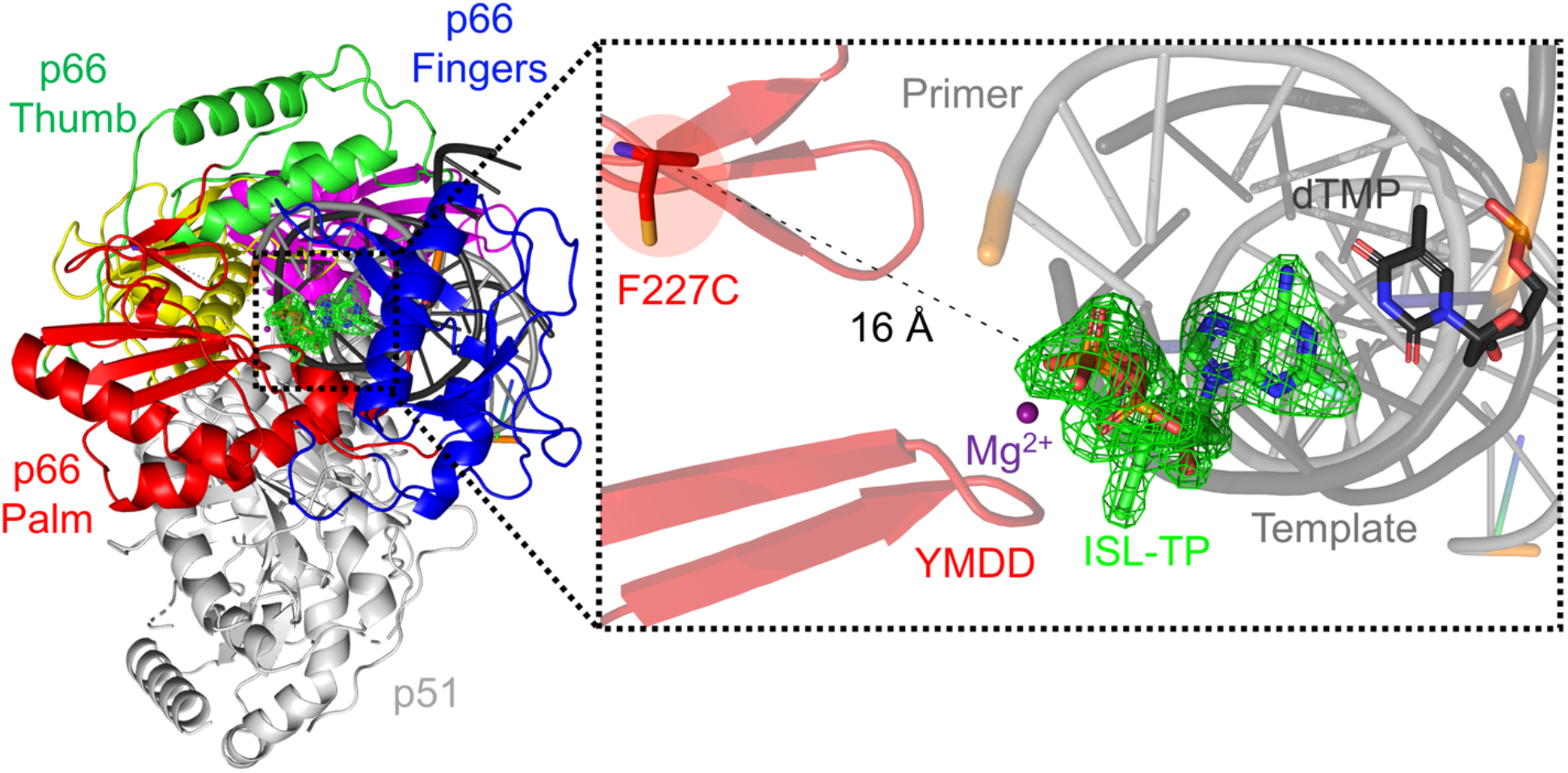
X-ray crystal structure of RT_F227C_/DNA_ddGMP_/ISL-TP complex at 1.8 Å. The p51 subunit is shown as a white cartoon, and the p66 subunit is shown in a cartoon as follows: fingers (blue), palm (red), thumb (green), connection (yellow) subdomains, and the RNase H domain (purple). Primer and template DNA are shown as light gray and dark gray cartoon, respectively. ISL-TP is shown in green sticks and colored by atom, binds at the polymerase active site of HIV-1 RT, ∼16 Å away from the F227C mutation at the NNRTI binding pocket. An Fo-Fc omit map is shown in green mesh contoured around ISL-TP at σ = 5.

### A 1.8 Å structure of F227C RT reveals propagated conformational changes linking the NNIBP to the polymerase active site and the excision channel

To visualize how an NNRTI-resistance mutation that is located >16 Å from the dNTP-binding site could enhance ISL activity, we determined the 1.8 Å crystal structure of F227C RT crosslinked to a ddGMP-terminated primer/template in complex with incoming ISL-TP (RT_F227C_/DNA_ddGMP_/ISL-TP, PDB:XXXX) (Figure 3, Supplemental Table 1). The structure shows clear electron density for ISL-TP within the polymerase active site and allows direct comparison with our previously determined corresponding complex with WT RT (RT_F227C_/DNA_ddGMP_/ISL-TP, PDB: 5J2M). Structural overlay of the WT and F227C complexes revealed that the F227C substitution triggers conformational rearrangements that propagate from the NNRTI-binding pocket toward the polymerase active site and excision machinery (Figure 4A).

**Figure 4.**
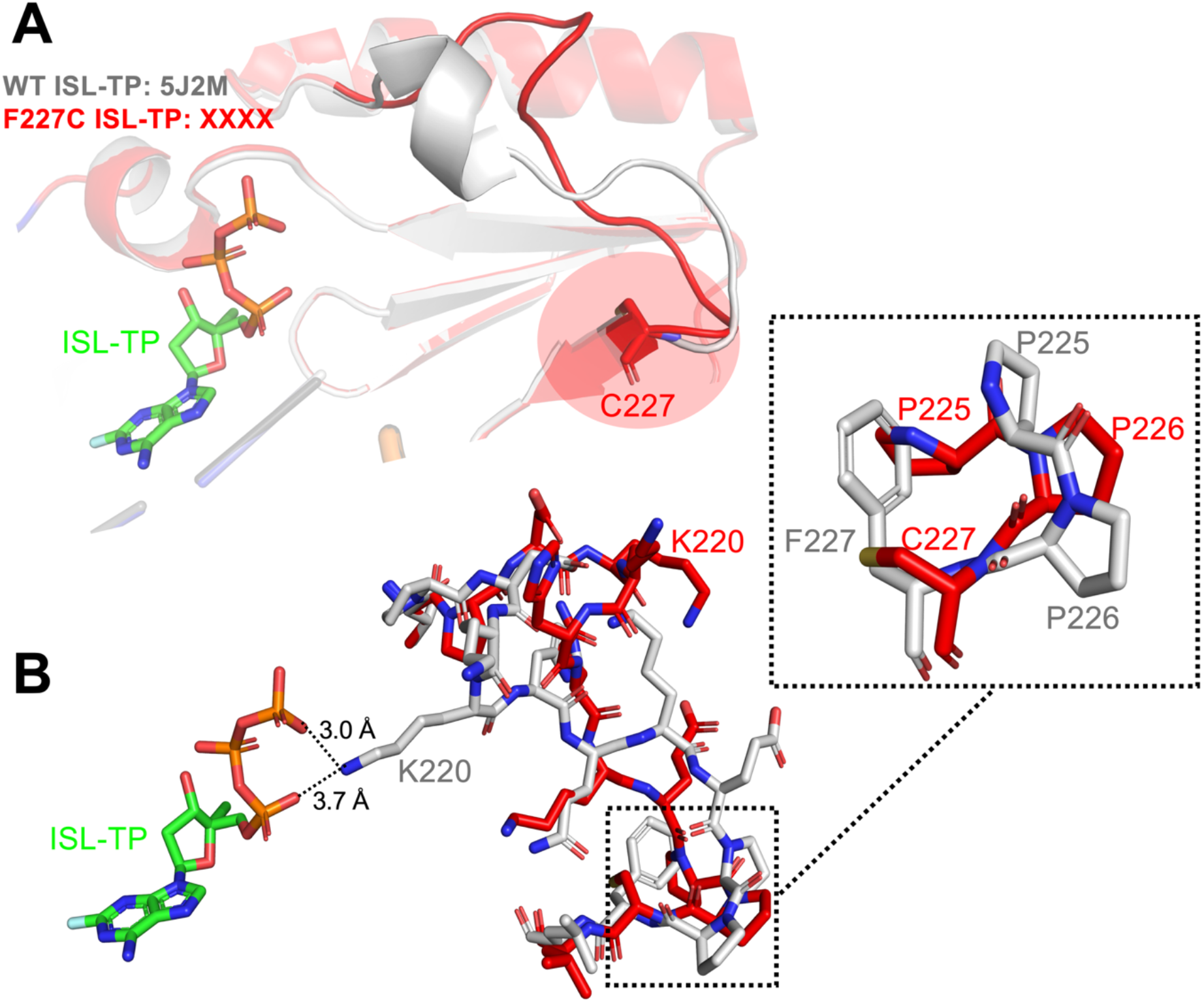
F227C induces conformational changes in regions associated with NNRTI binding and ATP-mediated excision. (A) Overlay of WT HIV-1 RT/dsDNA_ddGMP_/ISL-TP (gray; PDB: 5J2M) and F227C HIV-1 RT/dsDNA_ddGMP_/ISL-TP (red; PDB: XXXX) structures highlighting conformational changes induced by the F227C mutation. ISL-TP is shown in green sticks. The F227C mutation alters the positioning of structural elements proximal to the NNRTI-binding pocket and regions implicated in ATP-mediated excision. (B) Stick representation of residues surrounding positions 225–227 demonstrating local structural rearrangements caused by the F227C mutation. The inset highlights a shift in the backbone and side-chain orientations of residues P225–P227 that may contribute to altered inhibitor susceptibility and excision activity. Distances shown by a dotted black line.

The most pronounced structural differences were observed within residues 217-227, a region positioned at the base of the thumb subdomain and adjacent to the NNRTI-binding pocket. In WT RT, the bulky side chain of F227 stabilizes the local loop architecture through hydrophobic packing with P225, Q222, and neighboring residues. Q222 also makes H-binding interactions with the backbone of L228. Replacement of phenylalanine with the smaller cysteine allows the neighboring P225 to shift into the vacated space, destabilizing the 225-227 loop and causing an outward displacement of the backbone (Figure 4B). These local changes extend toward regions implicated in translocation dynamics and ATP-mediated excision. They reposition K220, open the nucleotide-binding region, and induce changes into the ATP-binding/excision channel.

### F227C modestly alters nucleotide incorporation

Increased susceptibility to a nucleoside RT translocation inhibitor (NRTTI) can arise through several distinct mechanisms, including a) altered nucleotide incorporation efficiency, b) impaired translocation, or c) defects in excision-based rescue pathways (46). To determine whether the F227C substitution affects ISL-TP incorporation, we performed steady-state single-nucleotide incorporation assays using both the natural substrate dATP and ISL-TP (Figure 5A, Supplemental Figure 2). Kinetic analysis demonstrated non-statistically significant changes (Welch’s t-test) in substrate utilization between WT and F227C RT. For dATP incorporation, a modest overall increase in the catalytic efficiency (k_cat_/K_m_) was observed (1.25-fold; Figure 6A). Thus, the overall differences between ISL-TP *vs*. dATP incorporation by WT and F227C RT cannot account for the pronounced ISL hypersusceptibility observed in biochemical and virological assays. Structural comparisons confirmed some remodeling around the polymerase active site, including repositioning of residue K220. In the WT enzyme, K220 is positioned closer to the incoming nucleotide-binding region, forming direct interactions with the triphosphate (Figure 4, 5), whereas in the F227C structure, this interaction network appears disrupted, producing a more open active-site architecture. These changes may increase accessibility of incoming nucleotides to the active site and subtly alter substrate positioning during catalysis. However, they may not significantly affect sensitivity to ISL-TP, as they are likely to affect similarly the incorporation of both inhibitor and natural triphosphate substrate. Of note, it is possible that active site interactions may vary with nucleic acid sequences; hence, a contribution of altered binding to ISL susceptibility cannot be entirely dismissed.

**Figure 5.**
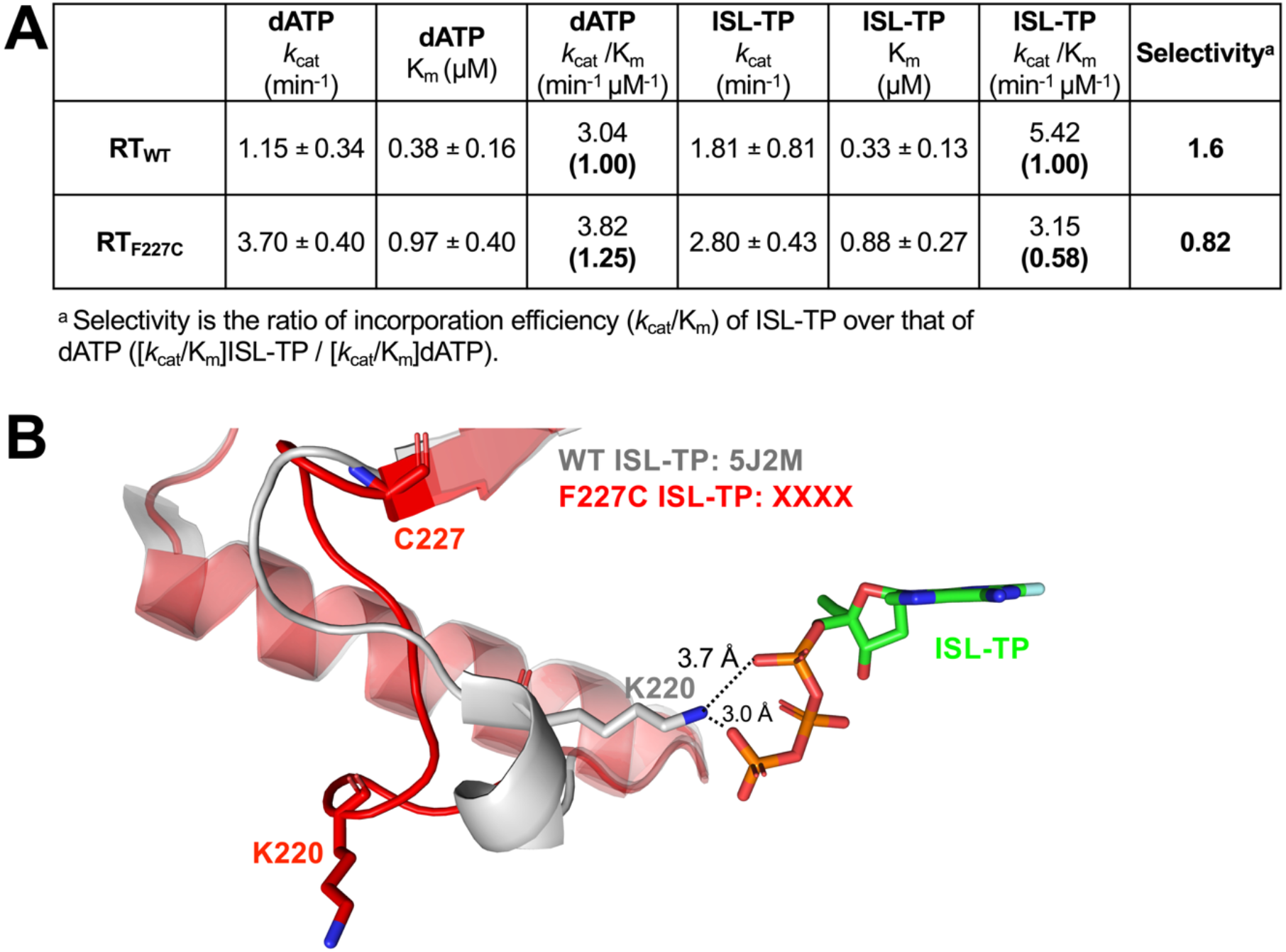
Biochemical and structural characterization of the effects of F227C on nucleotide incorporation. (A) Steady-state kinetic parameters for ISL-TP and dATP incorporation by WT and F227C HIV-1 RT (n = 3-4). (B) Overlay of WT HIV-1 RT/dsDNA_ddGMP_/ISL-TP (gray; PDB: 5J2M) and F227C HIV-1 RT/dsDNA_ddGMP_/ISL-TP (red; PDB: XXXX) structures illustrating how the F227C substitution alters the architecture of the polymerase active site, increasing active site accessibility for incoming dNTPs and significantly altering the positioning of K220.

**Figure 6.**
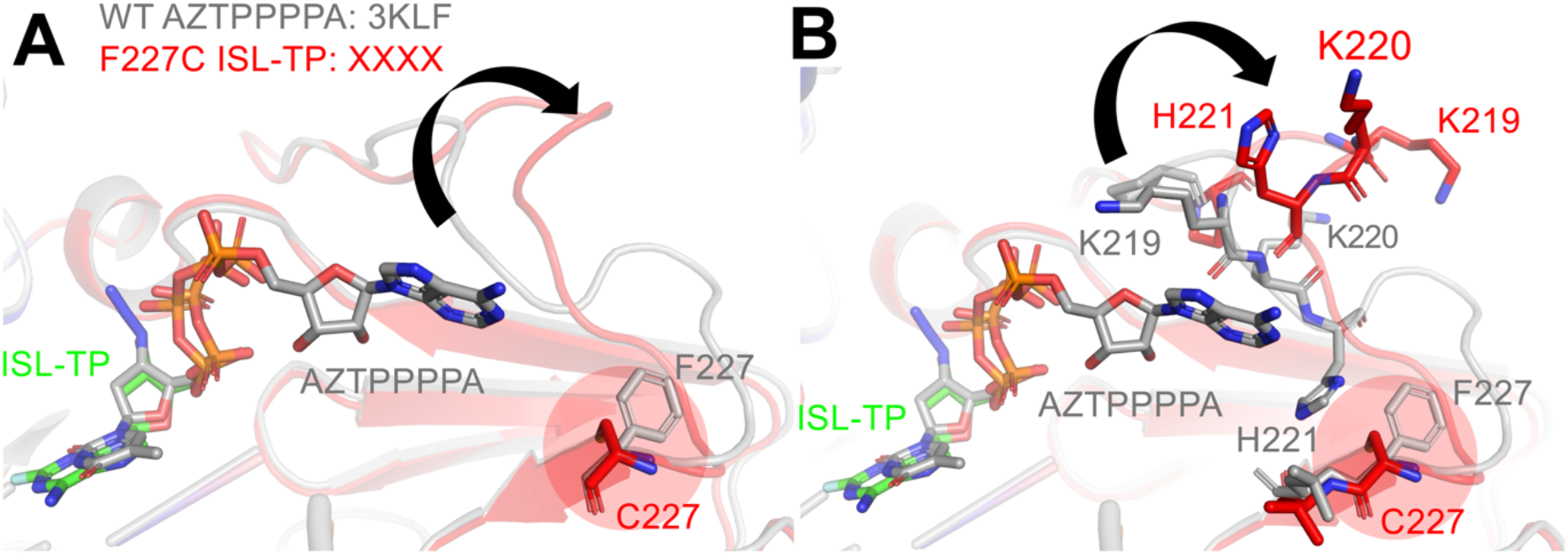
F227C restructures the ATP excision channel and impairs ATP-mediated rescue. A) Structural overlay of WT HIV-1 RT/dsDNA_ddGMP_ in complex with excision product AZTPPPPA (gray; PDB: 3KLF) and F227C HIV-1 RT/dsDNA_ddGMP_/ISL-TP (red; PDB: XXXX), showing conformational changes induced by the F227C mutation near the proposed ATP-binding/excision channel. ISL-TP is shown as green sticks, and AZT-PPPPA is shown as gray sticks. B) Enlarged view of residues surrounding positions 217–221 demonstrating rearrangements in the positioning of K219, K220, and H221 caused by the F227C mutation. These residues have previously been implicated in ATP-mediated excision activity.

### F227C profoundly impairs ATP-dependent excision

To further investigate the mechanism underlying F227C hypersusceptibility to ISL, we examined whether the substitution altered the ATP-mediated unblocking activities in the absence of DNA polymerization. Although ATP-dependent excision by WT RT is relatively low compared with RT bearing AZT resistance mutations forming a distinct ATP-binding pocket, we have previously shown that even WT RT can bind ATP for low-level excision, thereby affecting nucleoside analog susceptibility (47). Superposition of the relevant NRTI excision product (WT RT/nucleic acid/AZTppppA; PDB: 3KLF) (34) on the RT_F227C_/DNA_ddGMP_/ISL-TP structure (Figure 6A) revealed that the F227C substitution induces conformational rearrangements extending toward the ATP-binding channel. Most notably, the loop encompassing residues 217-221 appeared displaced outward relative to the WT structure, resulting in a more open and repositioned architecture near the excision site. Closer inspection of this region demonstrated altered positioning of residues K219, K220, and H221 in the F227C structure (Figure 6). These residues have been implicated in ATP-mediated excision activity (47), likely through ATP binding and positioning during “unblocking” of an incorporated nucleoside. Thus, we hypothesized that these structural rearrangements perturb the organization of the excision channel and affect ATP-based excision, despite being spatially distant from the polymerase active site itself. To address this hypothesis, we measured the effect of the F227C mutation on ATP-based rescue of ISL-MP-terminated primers at physiologically relevant ATP concentrations (3.5 mM) (48, 49). Notably, compared to WT RT, F227C RT had markedly impaired ATP-dependent ability to rescue ISL-terminated primers (Figure 7A). Michaelis–Menten analysis revealed that F227C primarily affected the apparent *k*_cat_, and to a lesser extent the apparent K_m_, causing an overall ∼12-fold reduction in the catalytic efficiency (*k*_cat_/K_m_) of ATP-dependent rescue, which is essentially the same *in magnitude* as the observed ISL hypersusceptibility of F227C (Figure 7B, C). Combined with the structural data, the results support the hypothesis that F227C distorts the ATP-binding/excision channel by repositioning the 217-221 loop and altering the orientation of K219-K221. These structural changes appear to contribute significantly to the pronounced ISL hypersusceptibility phenotype.

**Figure 7.**
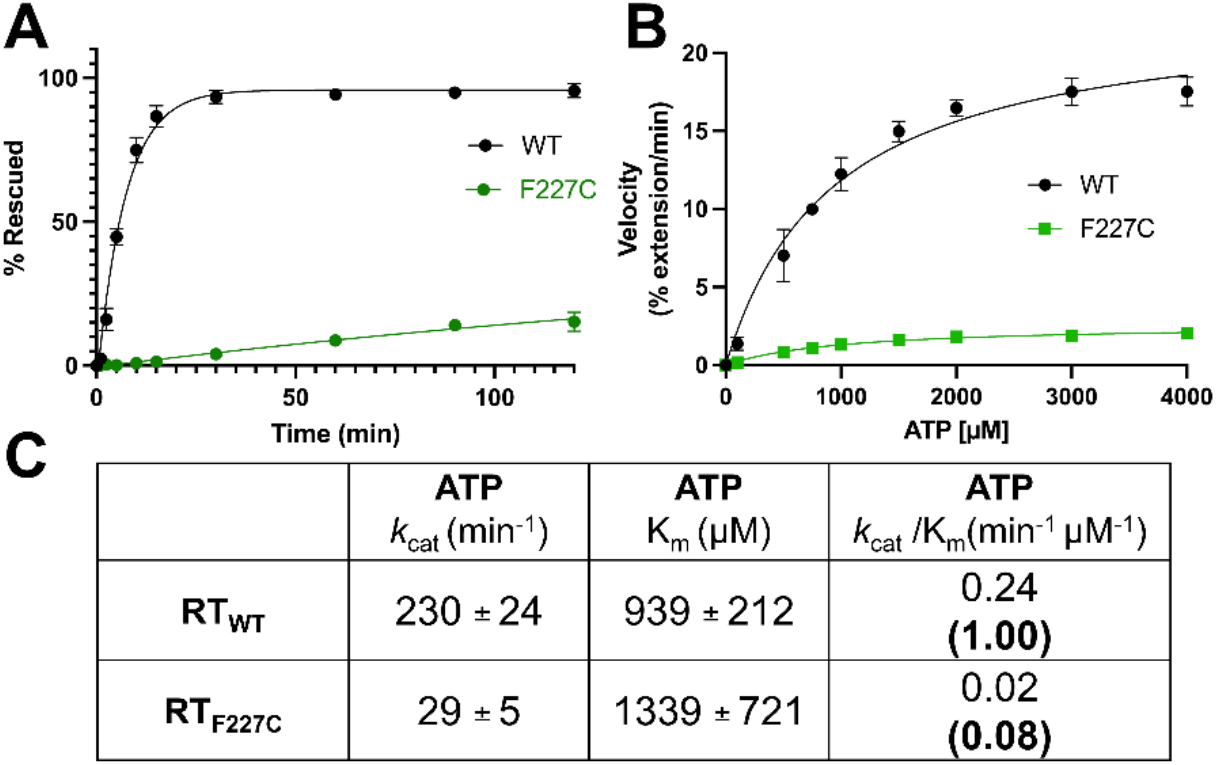
Effects of F227C on ATP-mediated excision efficiency. (A) Time-course analysis of ATP-mediated ISL-TP excision and rescue by RT_WT_ and RT_F227C_. The percent rescued product was plotted over time. (B) ATP-dependent excision and rescue velocities plotted as a function of ATP concentration (0–4000 μM ATP). Curves were fit using Michaelis–Menten kinetics in GraphPad Prism. (C) Steady-state Michaelis–Menten analysis of ATP-mediated *excision and rescue reactions to determine apparent k*_cat_ and K_m_ values. Relative catalytic efficiencies normalized to RT_WT_ are shown in parentheses. Data represent mean ± SD from at least two independent experiments.

### F227C reduces rather than enhances the translocation of ISL-terminated primers

Hypersusceptibility to translocation inhibitors can theoretically arise from enhanced forward translocation, thereby protecting against excision. The translocation state of WT and F227C RT complexes was evaluated using Fe^2+^ hydroxyl radical footprinting assays on ISL-MP-terminated complexes. These assays were conducted at increasing concentrations of the next incoming nucleotide, dTTP, added to promote formation of the post-translocated complex (Supplemental Figure 4). Consistent with previous studies, cleavage at position −18 corresponded to the pre-translocated state, whereas cleavage at position −17 indicated formation of the post-translocated complex. Quantification of post-translocated complexes showed that WT RT more readily transitioned into the post-translocation state following ISL-MP incorporation. In contrast, F227C RT exhibited decreased formation of the post-translocated state with increased retention in the pre-translocated register (Figure 8). The reduced translocation efficiency is consistent with the primer extension data, which demonstrate enhanced stabilization of ISL at immediate chain-termination sites in the mutant enzyme. This further supports impaired excision, rather than accelerated translocation, as the principal mechanism of ISL hypersusceptibility.

**Figure 8.**
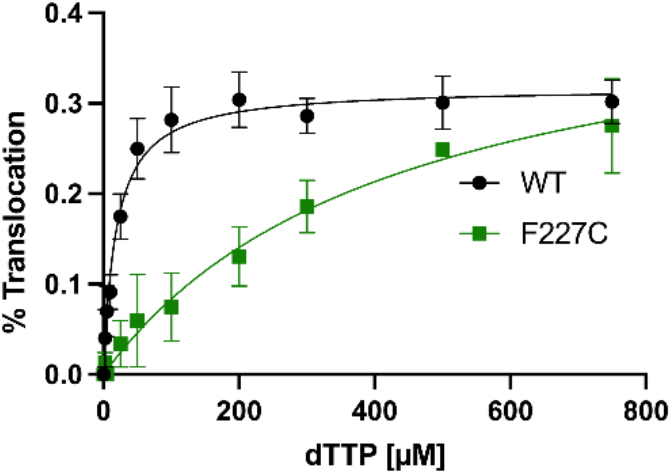
Effect of F227C mutation on the translocation state of RT bound to T/P_ISL-MP_. The translocation state of HIV-1 RT after ISL-MP incorporation was determined using site-specific Fe^2+^ footprinting. T_d43_/P_d30-ISL-MP_ (100 nM) with 5’-Cy3-label on the DNA template was incubated with WT or F227C HIV-1 RT (600 nM) and various concentrations of the next incoming nucleotide (dTTP). The complexes were treated for 5 minutes with ammonium iron sulphate (1 mM) and resolved on a polyacrylamide 7 M urea gel. Excision at position −18 indicates a pre-translocation complex, while excision at position −17 represents a post-translocation complex. The post-translocated complexes were determined from the gels and plotted using GraphPad Prism (n=3).

### ISL and DOR retain additive antiviral activity against F227C despite high-level DOR resistance

Finally, we asked whether combining ISL with NNRTIs could be clinically beneficial in the F227C background. The ISL hypersusceptibility of F227C HIV-1 could be therapeutically leveraged in combination with NNRTIs. We evaluated the antiviral interactions between ISL and DOR or ESV against WT and F227C HIV-1 using multidimensional synergy analysis (Figure 9, Supplemental Figure 5, Supplemental Figure 6C, D). TDF was also tested in combination with DOR against WT and F227C HIV-1 (Supplemental Fig 6 A, B, D). Inhibition from combination dose-response matrices was analyzed using the HSA, Bliss, Loewe, and ZIP reference models in SynergyFinder Plus (50) to assess whether the inhibitors exhibited synergistic, additive, or antagonistic effects.

**Figure 9.**
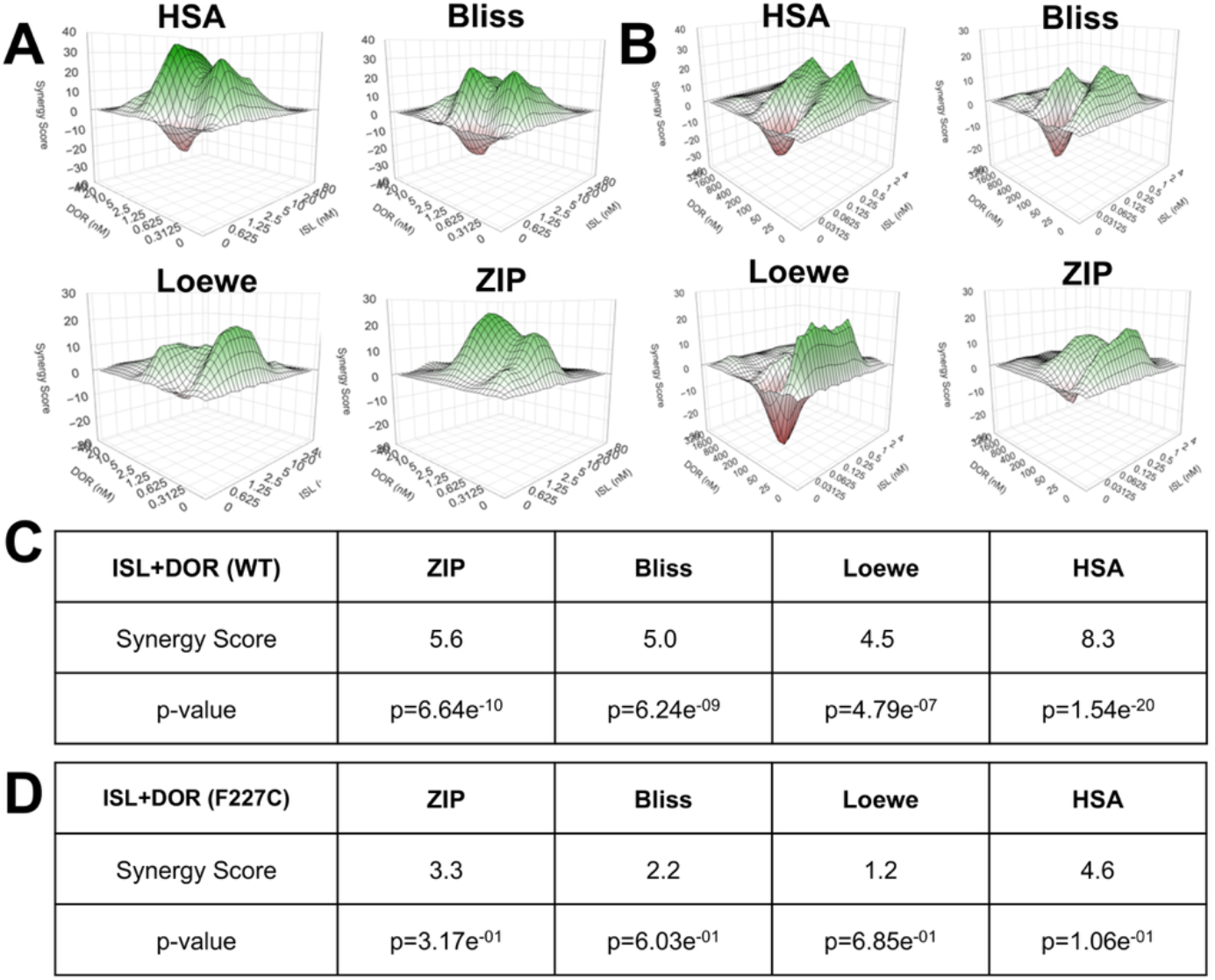
Synergy analysis of ISL and DOR combinations against WT and F227C HIV-1. Dose-response matrices and three-dimensional synergy landscapes for ISL and DOR combinations against WT (A) and F227C (B) pNL4-3 Δ env VSVG pseudotyped virus were generated using the HSA, Bliss, Loewe, and ZIP reference models from SynergyFinder Plus outputs. Graphs were generated with R (version 4.5.0). Green regions indicate synergistic interactions, whereas red regions indicate antagonistic interactions. Average synergy scores and associated p-values for each model are shown below the corresponding plots (WT-C, F227C-D). Synergy scores between −10 and 10 are interpreted as additive effects, while scores >10 and <−10 indicate synergistic and antagonistic interactions, respectively. Overall, ISL and DOR demonstrated predominantly additive synergistic interactions against both WT and F227C HIV-1 (n=3).

Across both WT and F227C viruses, the ISL and DOR combination demonstrated predominantly additive interactions, with average synergy scores ranging from −10 to 10 for all models tested (Figure 9C, D). The WT virus produced average synergy scores ranging from 4.5 to 8.3, depending on the scoring model, whereas F227C values ranged from 1.2 to 4.6. Combination therapy of ISL + ESV against the F227C virus, had synergy scores ranging from -2.2 to 1.1 (Supplemental Figure 6C, D), also suggesting this combination is additive. These findings indicate that the ISL/DOR and ISL/ESV combinations remain broadly additive despite the strong resistance of F227C to DOR and ESV alone.

Although the overall interaction profile was classified as additive, localized regions of enhanced synergy were observed at specific inhibitor concentrations within the three-dimensional synergy landscapes (Figure 9A, B, Supplemental Figure 6A-C). These concentration-dependent synergistic regions suggest that under certain inhibitor ratios, simultaneous targeting of RT by both ISL and DOR may produce enhanced suppression of viral replication beyond what would be expected from either inhibitor individually. Importantly, no strong antagonistic interactions were observed across the tested concentration ranges.

Notably, despite the substantial resistance of F227C to DOR monotherapy, the ISL/DOR and ISL/ESV combinations maintained additive antiviral activity against the mutant virus. These findings suggest that ISL-containing regimens may partially preserve antiviral efficacy even in the presence of NNRTI resistance mutations such as F227C and support further investigation of ISL-based combination strategies targeting structurally and mechanistically distinct sites within HIV-1 RT. The combination of TDF and DOR produced additive synergy scores ranging from 0.016 to 3.6 in WT virus (Supplemental Figure 6A, D) and from −2.8 to 1.7 in F227C virus (Supplemental Figure 6B, D). This combination was evaluated because TDF, like ISL, is a nucleos(t)ide analog targeting RT and is widely used as part of standard-of-care antiretroviral therapy (51). These findings indicate that TDF and DOR exhibit primarily additive antiviral activity, although the overall synergy was lower than that observed for the ISL and DOR combination.

## Discussion

With the recent FDA approval of ISL in combination with DOR (16), ISL is poised to play an important role in future HIV-1 treatment strategies. ISL is a highly potent picomolar inhibitor of HIV; it exhibits a high barrier to resistance and has strong potential as a long-acting therapeutic, a key goal for next-generation antiretroviral therapies (18, 52, 53) (54). Hypersusceptibility occurs when a specific drug-resistance mutation makes HIV-1 *more* vulnerable to antiretrovirals. It is well-documented in antiviral therapies, between NRTIs and occasionally between NRTI and NNRTIs (48). For example, the NRTI resistance mutation M184V/I was reported to cause hypersusceptibility to NRTIs AZT and TDF (22); similarly, TDF-resistance K65R causes hypersusceptibility to AZT (22). NRTI-resistance mutations M41L, M184V, L210W, and T215Y or M41L/T215Y and M41L/T215Y/M184V were associated with virological suppression for NNRTI (efavirenz)-treated patients (49-52); Therefore, drug hypersusceptibility may create opportunities for “resistance-informed” combination regimens that exploit these vulnerabilities.

Here we determined the biochemical and structural basis underlying the unexpected hypersusceptibility of the NNRTI resistance mutation F227C to the NRTTI ISL. In virological assays we initially confirmed the hypersusceptibility phenotype of F227C, showing that this RT mutation confers 11.9-fold hypersusceptibility to ISL (Figure 2) (37, 44). In contrast, F227C did not increase susceptibility to tenofovir, another adenosine analog within the NRTI class. This difference may reflect the distinct excision properties of TDF-terminated primers, as tenofovir is not efficiently excised by ATP unless thymidine analog mutations (TAMs) are present (55). We additionally observed high-level resistance to DOR and ESV (Figure 2), but only moderate resistance to RPV and ETR. These differences likely arise from distinct inhibitor-binding modes within the NNRTI-binding pocket. Both DOR and depulfavirine, the active form of pro-drug ESV, make direct interactions with F227 (42, 56), whereas RPV and ETR adopt U-shaped flexible conformations that may rely less heavily on interactions with this residue (57, 58). However, it remains unclear how the F227C substitution alters the architecture of the NNRTI-binding pocket in the presence of bound NNRTIs or NRTTIs.

Biochemical analysis further demonstrated that F227C-associated hypersusceptibility is dependent on ATP-containing conditions (Figure 2, 7). In the absence of ATP, hypersusceptibility was not observed, similar to prior findings reported for K65R RT and ISL (36). These results suggest that post-incorporation events, rather than incorporation alone, play a dominant role in the phenotype. Primer extension assays revealed altered stopping patterns and changes in translocation behavior, particularly at the P+6/P+7 positions following ISL incorporation. Our previous studies have demonstrated that translocation states are sequence dependent (45); however, to our knowledge, this is the first observation of a mutant RT directly altering ISL-associated translocation behavior. ATP also altered polymerization patterns, potentially through allosteric effects on RT and dNTP incorporation (59), as reflected by distinct pausing and extension profiles observed in the presence of ATP.

Our structural studies provide a mechanistic explanation for these biochemical observations. Although F227 is positioned >16 Å away from the polymerase nucleotide binding site (N-site), the F227C substitution induced substantial long-range conformational rearrangements extending beyond the immediate NNRTI-binding pocket. Replacement of the bulky aromatic phenylalanine with cysteine disrupted local packing interactions within the 217-227 loop region and increased conformational flexibility at the base of the thumb subdomain (Figure 4). These perturbations appear to propagate through adjacent structural elements, with 227 serving as the nexus of these changes that alter the positioning of the 217-227 loop. Notably, residues K219, K220, and H221 in this loop have all been previously implicated in either RT catalytic dynamics or excision activity (47). Structural comparison with WT RT complexes further revealed outward displacement of the 217-221 loop near the proposed ATP-binding/excision channel, suggesting that F227C reorganizes the excision machinery despite being spatially distal from the active site itself.

Steady-state kinetic analyses demonstrated only modest differences in incorporation efficiency between WT and F227C RT. WT RT displayed a small preference for ISL-TP over the endogenous substrate dATP, whereas F227C mildly favored dATP incorporation. Although previous studies have also reported modest increases in ISL incorporation efficiency by F227C RT (43), differences in experimental design, including pre-steady-state *versus* steady-state methodologies, and different template sequence context, confound direct comparisons. While altered incorporation kinetics may contribute partially to hypersusceptibility, our data suggest they are unlikely to fully account for the magnitude of the phenotype. Structurally, we also observed widening of the dNTP entrance channel and altered positioning of K220, consistent with subtle changes in substrate accessibility and active-site organization (Figure 5).

The most substantial functional defects associated with F227C were observed during excision and rescue assays (Figure 6, 7). F227C RT exhibited impaired rescue of ISL-MP-terminated primers with ATP, with substantially greater impairment observed during ATP-dependent rescue. To determine whether this defect resulted from altered substrate binding or impaired excision efficiency, we performed steady-state kinetic analyses. Only modest differences in apparent K_m_ values were observed between WT and F227C RTs for ATP, whereas the mutant enzyme displayed markedly reduced apparent *k*_cat_ values in ATP excision. These findings suggest that F227C primarily impairs the catalytic step of the excision reaction rather than substrate recognition. Notably, ATP is not expected to bind tightly under these conditions (47, 59), further supporting a catalytic defect based on misalignment of bound ATP rather than a major binding deficiency. Additionally, it should be emphasized that this assay does not directly measure substrate binding or excision alone, as the observed kinetics result from a combination of excision and rescue (subsequent primer extension events). Therefore, more direct biochemical and biophysical approaches may be required to accurately determine substrate binding affinities and single-turnover excision kinetics. However, based on our incorporation studies measuring the catalytic efficiency of dNTP incorporation by WT and F227C RT, we find no evidence of a major global enzymatic defect associated with the F227C substitution. Together, these data indicate that the reduced rescue activity of F227C RT primarily results from impaired excision efficiency rather than defects in rescue. Hence, ISL-MP may remain blocked at the primer terminus for longer periods in the mutant enzyme, potentially contributing to enhanced inhibition of DNA synthesis. This interpretation is also consistent with the structural displacement of the proposed ATP-binding/excision loop observed in the F227C structure (Figure 6). Importantly, these defects are unlikely to substantially affect excision of endogenous nucleotides, as natural dNMPs are not readily excised by RT (55).

In parallel, F227C reduced forward translocation on primers terminated with ISL-MP. If hypersusceptibility was due to increased translocation, increased movement into the post-translocated P-site should protect ISL-MP from excision (Figure 8). Instead, the observed decrease in translocation argues against this mechanism and further supports defective excision as the primary contributor to hypersusceptibility.

Collectively, our data support a model in which F227C reshapes the conformational landscape of HIV-1 RT, indirectly enhancing ISL sensitivity through disruption of ATP-mediated rescue pathways. Although modest effects on incorporation kinetics and translocation were observed, impaired excision appears to represent the dominant mechanistic driver of hypersusceptibility. These findings further emphasize the extensive conformational coupling among the NNRTI-binding pocket, the polymerase active site, translocation-associated regions, and the ATP-mediated excision machinery within HIV-1 RT.

The complementary resistance profiles of DOR or ESV with ISL also suggest potential therapeutic advantages for these combinations (60, 61). While F227C confers resistance to DOR and ESV, it simultaneously increases susceptibility to ISL, potentially creating an evolutionary tradeoff that may suppress the emergence of resistant viral populations. Consistent with this hypothesis, synergy analyses demonstrated predominantly additive interactions between ISL and DOR in both WT and F227C viral backgrounds (Figure 9). These data are consistent with other studies recently published (62). These findings may help guide optimization of antiviral dosing strategies while potentially reducing overall drug burden. More broadly, combinations of NRTTIs and NNRTIs may be particularly effective due to their ability to promote highly inhibited or “dead-end” RT complexes (63). ISL stabilizes translocation-defective states following nucleotide incorporation, while DOR restricts conformational flexibility through binding within the NNRTI-binding pocket. Together, these inhibitors may cooperatively impair nucleotide incorporation, translocation, and excision-mediated rescue, thereby reducing the ability of RT to recover productive DNA synthesis.

Overall, this work demonstrates that resistance-associated mutations can create previously unrecognized mechanistic vulnerabilities within the HIV-1 RT. Rather than functioning solely as determinants of inhibitor escape, mutations such as F227C can reshape RT conformational dynamics, thereby sensitizing the enzyme to mechanistically distinct inhibitors. These findings provide new insight into the structural coupling that governs RT function and may inform the future development of combination therapies that exploit hypersusceptibility pathways in HIV-1.

## Materials and Methods

### Reverse Transcriptase Expression and Purification

#### RT construct for enzymology-based assays

HIV-1 RT (p6HRT-prot) was expressed in JM109 cells (Agilent Technologies, Inc.) and purified by nickel affinity chromatography (Cytiva) as previously described (64). The F227C substitution was introduced into the RT coding sequence using the Q5® Site-Directed Mutagenesis Kit (New England Biolabs).

#### RT construct for Crystallography-based studies

BL21 (DE3)-RIL cells (Agilent Technologies, Inc.) were transformed with a dual-vector system containing the p66 and p51 RT subunits (65). RT was expressed and purified by nickel affinity chromatography and MonoQ anion-exchange chromatography, as previously described. The F227C substitution was introduced into the RT coding sequence using the Q5® Site-Directed Mutagenesis Kit (New England Biolabs) and confirmed by sequencing (Genewiz).

Purified RT (65) was crosslinked with dsDNA (primer sequence: 5′-ACA GTC CCT GTT CGG G*CG CC-3′ (G* has a thioalkyl tether covalently attached to the N^2^ group of the guanosine) and template sequence: 5′-ATG GTC GGC GCC CGA ACA GGG ACT GTG-3′; 5′-ATG GAT GGC GCC CGA ACA GGG ACT GTG-3′) (Trilink Technologies and Integrated DNA Technologies). dsDNA was prepared by mixing the template primer in a 1:1 ratio (0.1 mM), heating at 95 °C for 5 minutes, and then allowing it to cool at room temperature for 30 minutes. 15 μM annealed template/primer (T/P) substrates were incubated with 20 μM RT in 25 mM Tris-HCL, pH 8.0, 10 mM MgCl_2_, and 0.1 mM 2′,3′-dideoxyguanosine-5′-triphosphate (ddGTP) (AAT Bioquest), 25 mM Tris, pH 8.0, and 100 mM NaCl overnight at 30 °C to obtain the RT/DNA_ddGMP_ complex. Cross-linked RT-dsDNA complex was purified using a Ni-Heparin tandem column strategy (66) and evaluated using non-reducing SDS-PAGE (Novex). Fractions containing DNA–cross-linked RT were pooled and concentrated to 15 mg/mL. To obtain the RT/DNA_ddGMP_/ISL-TP ternary complex, the initial RT/DNA_ddGMP_ complex sample was incubated at room temperature for 15 min with 1.5 molar excess of ISL-TP (MedChemExpress) and 5 mM MgCl_2_. Hanging drops were prepared by mixing 1:1, 2:1, or 2:1 volume ratio of the well solution and the ternary complex. Crystals were grown at 18 °C in 7-9% (wt/vol) PEG 8000, 25 mM bis-tris propane (AmBeed, Inc.), pH 6.0, and 5 mM magnesium chloride. Crystals were cryoprotected by elevating PEG 4000 (Hampton Research Corp.) up to 35% (wt/vol) over 1 h and were flash cooled in liquid nitrogen.

#### X-ray Data Collection and Determination

X-ray diffraction data were collected on a DECTRIS Eiger 16 M detector at the SouthEast Regional Collaborative Access Team (SERCAT) beamline 22-ID of the Advanced Photon Source (APS) at Argonne National Laboratory. Datasets were processed using XDS (67) and indexed in space group P2_1_ (a=83.99, b=100.68, c=92.49 Å; β=113.95°) with one RT per asymmetric unit. The crystal structure was solved by molecular replacement using PDB ID 5J2M (66) as starting model in PHASER (68) in the CCP4 package (69). Several rounds of manual rebuilding using Coot (70) and refinement with PHENIX (71) and Refmac5 (72) were performed. Final model validation was performed using MolProbity (73) and the wwPDB validation server. Data collection, processing, and refinement statistics are available in Table S1. The structure has been deposited in the PDB, but the coordinates are on hold.

### Pseudotyped Viruses and Cell Culture Experiments

DMEM/High Glucose (Cytiva HyClone− Dulbecco’s Modified Eagle’s Medium cat no. SH30243.02) with 10% FBS, and 50 U/mL penicillin/50µg/mL streptomycin (Thermo Fisher) was used to culture HEK293T/17 (74) and TZM-green fluorescent protein (GFP) (Massimo Pizzato) (75) cell lines.

#### Pseudotyped virus production and titer

To produce the F227C, NL4-3Δenv plasmids, single-nucleotide mutagenesis was performed in the RT sequence using the NEB HiFi Assembly Kit (New England Biolabs, 2022). The plasmid was then sequenced to verify the presence of each mutation (Genewiz). To prepare WT and mutant VSVG-pseudotyped virus stocks, 5 µg of pNL4-3Δenv plasmids were each transfected with 1.5 µg of pVSVG in HEK293T cells using Xtreme-GENE HP transfection reagent (Roche, Switzerland). 48 h post-transfection, supernatant was collected, and infectious unit counts were determined for the NL4-3Δenv mutant and WT pseudotyped virus stocks in TZM-GFP cell lines.

#### Dose Response Curves and Synergy

For synergy experiments and dose response experiments, serially diluted ISL (Life Chemicals, Burlington, Ontario, Canada) and DOR (MedChemExpress, Cat. No.: HY-16767), ESV (Viriom, Inc.), RPV (Bei Resources HRP:12147), and ETR (Bei Resources HRP-11609) were plated in 96-well plates together with TZM-GFP cells. For synergy experiments, drugs were plated in a matrix format, 2- to 5-fold dilutions. Cells plus inhibitor were then incubated for 24 h. After 24 h, the cells were infected with a mixture of DEAE-dextran (1 µg/mL final concentration) and NL4-3Δenv WT or F227C VSVG pseudotyped virus. After infection, the plates were incubated for an additional 48 h. A Cytation 5 (BioTek, Winooski, VT, USA) was used to image GFP-positive cells, which were subsequently counted using Gen5 v3.15 (BioTek). To determine percent inhibition, technical replicates were averaged, and dose-response data were normalized to no-drug controls (Supplemental Figure 1, 3). Drug interaction analyses were performed using SynergyFinder Plus software (50) using the ZIP, Bliss, Loewe, and HSA reference models. Synergy scores between −10 and 10 were interpreted as additive, whereas scores >10 or <−10 were considered synergistic or antagonistic, respectively. Experiments were conducted in triplicate with technical duplicates.

### Biochemistry

#### Gel-based Primer Extension Assay

DNA template was annealed to a 5′-Cy3-labeled DNA primer (3:1 molar ratio; Td_31_/Pd_18_-P_0_). Primer extension assays were performed by incubating the DNA/DNA substrate (20 nM) with WT or F227C HIV-1 RT (20 nM) at 37°C in RT buffer containing 50 mM Tris-HCl (pH 7.8) and 50 mM NaCl. Reactions were initiated by the addition of ISL-TP and MgCl_2_ (final concentration of 6 or 10 mM) in a total reaction volume of 20 μL. All dNTPs were included at a final concentration of 1 μM in the presence or absence of 3.5 mM ATP. After 15 min, reactions were terminated with an equal volume of 100% formamide containing trace amounts of bromophenol blue. Reaction products were separated on 15% polyacrylamide/7 M urea gels and visualized using a Typhoon FLA 9000 PhosphorImager (GE Healthcare, NJ). Bands corresponding to fully extended products were quantified using Multi Gauge software. Data from at least 3 independent experiments were plotted as percent full extension, and IC_50_ values for ISL-TP were determined by one-site competition nonlinear regression analysis in GraphPad Prism 10.

##### Template: Td_**31**_

5′-CCA TAG CTA GCA TTG GTG CTC GAA CAG TGA C-3′

##### Primer: Pd_**18**_**-P**_**0**_

5′-Cy3-GTC ACT GTT CGA GCA CCA-3′

### Single Nucleotide Incorporation Assay under steady-state kinetics

#### Excision/Rescue Assay

Steady-state kinetic parameters (K_m_ and *k*_cat_) for ISL-TP or dATP (Thermo Scientific) incorporation were determined using gel-based single-nucleotide incorporation assays performed under saturating template/primer (T/P) conditions (10-fold excess over RT). Reactions contained RT buffer, 6 mM MgCl_2_, 100 nM Td_31_/Pd_18_-P_0_ substrate, 10 nM WT or F227C HIV-1 RT, and increasing concentrations (0–25 μM) of dATP or ISL-TP in a final reaction volume of 20 μL. Reactions were incubated at 37 °C for 1–5 min and terminated with an equal volume of 100% formamide containing trace bromophenol blue. Products were resolved on 15% polyacrylamide/7 M urea denaturing gels and visualized using a Typhoon FLA 9000 PhosphorImager (GE Healthcare). Band intensities corresponding to incorporated products were quantified using AzureSpot Pro software (Azure Biosystems). Initial velocities were calculated as the percentage of incorporated product formed over time and plotted as a function of substrate concentration. Kinetic parameters were determined by fitting the data to the Michaelis–Menten equation using GraphPad Prism 10. Experiments were performed in two to four independent replicates, and mean values ± standard deviations were calculated.

#### Excision Rescue Assay

Template/primer substrates terminated with ISL-monophosphate at the 3′ primer terminus (T/P_ISL-MP_) were generated by incubating 500 nM Td_31_/Pd_18_-P_0_ with 1 μM HIV-1 RT in RT buffer containing 10 mM MgCl_2_. ISL-TP was added, and reactions were incubated at 37 °C for 1 h to allow complete incorporation. T/P_ISL-MP_ products were purified using a QIAquick Nucleotide Removal Kit (Qiagen, Valencia, CA).

For ATP-dependent rescue assays, 20 nM purified T/P_ISL-MP_ substrate was incubated with 60 nM WT or F227C HIV-1 RT in RT buffer containing 10 mM MgCl_2_, 3.5 mM ATP (Thermo Scientific), 100 μM dATP, 0.5 μM dTTP, and 10 μM ddGTP. Reactions were carried out at 37 °C, and aliquots were collected at the indicated time points (0–90 min). Reaction products were resolved on 15% polyacrylamide/7 M urea denaturing gels and visualized using a Typhoon FLA 9000 PhosphorImager (GE Healthcare). Band intensities corresponding to rescued and extended products were quantified using AzureSpot Pro software (Azure Biosystems).

For ATP concentration-dependent excision/rescue kinetic assays, reactions were performed under similar conditions with increasing ATP concentrations while maintaining constant concentrations of RT, T/P_ISL-MP_ substrate, dATP, dTTP, and ddGTP. Initial velocities were determined from the formation of rescued products over time and plotted as a function of ATP concentration. Apparent kinetic parameters were determined by fitting the data to the Michaelis–Menten equation using GraphPad Prism 10. Experiments were performed in at least two independent replicates, and mean values ± standard deviations were calculated.

#### Hydroxyl Radical Footprinting Assay

Site-specific Fe^2+^ footprinting assays were performed using 5′-Cy3-labeled DNA templates. Briefly, 100 nM 5′-Cy3-Td_43_/Pd_30_ was incubated with 600 nM WT or F227C HIV-1 RT in a reaction buffer containing 120 mM sodium cacodylate (pH 7.0), 20 mM NaCl, 6 mM MgCl_2_, and 1 μM ISL-TP to achieve quantitative chain termination. Before Fe^2+^ treatment, complexes were pre-incubated for 7 min with increasing concentrations of the next incoming nucleotide (dTTP). Complexes were then treated with 1 mM ammonium iron sulfate as previously described (76) (77). This assay relies on Fe^2+^ autoxidation to generate localized hydroxyl radicals that cleave the nucleic acid near the RNase H active site, enabling analysis of RT translocation states (78). Experiments were performed in at least three independent replicates.

##### Primer: Pd_**30**_

5′-TCT ACA CTG ATT GTC ACT GTT CGA GCA CCA-3′

##### Template: Td_**43**_

5′-Cy3-CCA TAG ATA GCA TTG GTG CTC GAA CAG TGA CAA TCA GTG TAG A-3′

### Statistics

Statistical analyses were performed using GraphPad Prism 10 (GraphPad Software, San Diego, CA). EC_50_ and IC_50_ values were determined by nonlinear regression using a four-parameter dose-response model. Steady-state kinetic parameters were obtained by fitting data to the Michaelis– Menten equation. Statistical significance was evaluated using one-way ANOVA with Dunnett’s multiple-comparison test, one-sample t-tests, or Wilcoxon signed-rank tests as indicated in the figure legends. P values < 0.05 were considered statistically significant.

### Structural Analysis and Visualization

Structural alignments, overlays, and visualization of HIV-1 RT complexes were performed using PyMOL (Schrödinger LLC). Structural comparisons between WT and F227C RT complexes were generated by superposition of corresponding Cα atoms. Figures depicting conformational rearrangements, active-site architecture, and inhibitor interactions were prepared using PyMOL. Measurements and residue visualization were generated within the PyMOL molecular graphics system (79).

## Acknowledgments

A.A.S. acknowledges the Cold Spring Harbor Laboratory Macromolecular Crystallography Course for training and scientific support. We thank Massimo Pizzato for providing TZM-GFP cells.

X-ray data were collected at Southeast Regional Collaborative Access Team (SER-CAT) 22-ID beamline at the Advanced Photon Source, Argonne National Laboratory. SER-CAT is supported by its member institutions, equipment grants (S10_RR25528, S10_RR028976 and S10_OD027000) from the National Institutes of Health, and funding from the Georgia Research Alliance. This research used resources of the Advanced Photon Source, a U.S. Department of Energy (DOE) Office of Science user facility operated for the DOE Office of Science by Argonne National Laboratory under Contract No. DE-AC02-06CH11357.

## Author contributions

A.A.S., K.A.K., E.M., and S.G.S. conceived the project.

A.A.S., I.L.K., J.M., R.L.S., K.A.K., E.M., and S.G.S. designed the experiments.

A.A.S., I.L.K., and J.M. performed the experiments.

A.A.S., K.A.K., and S.G.S. analyzed the data, prepared the figures and supplementary materials, deposited the structures, and wrote the manuscript.

All authors reviewed and edited the manuscript.

## Data availability

All data needed to evaluate the conclusions in the paper are present in the paper and/or the Supplementary Materials.

