## Supplemental Figure 1 for "Structural and Mechanistic Basis of F227C-Mediated Hypersusceptibility to Islatravir in HIV-1 Reverse Transcriptase"

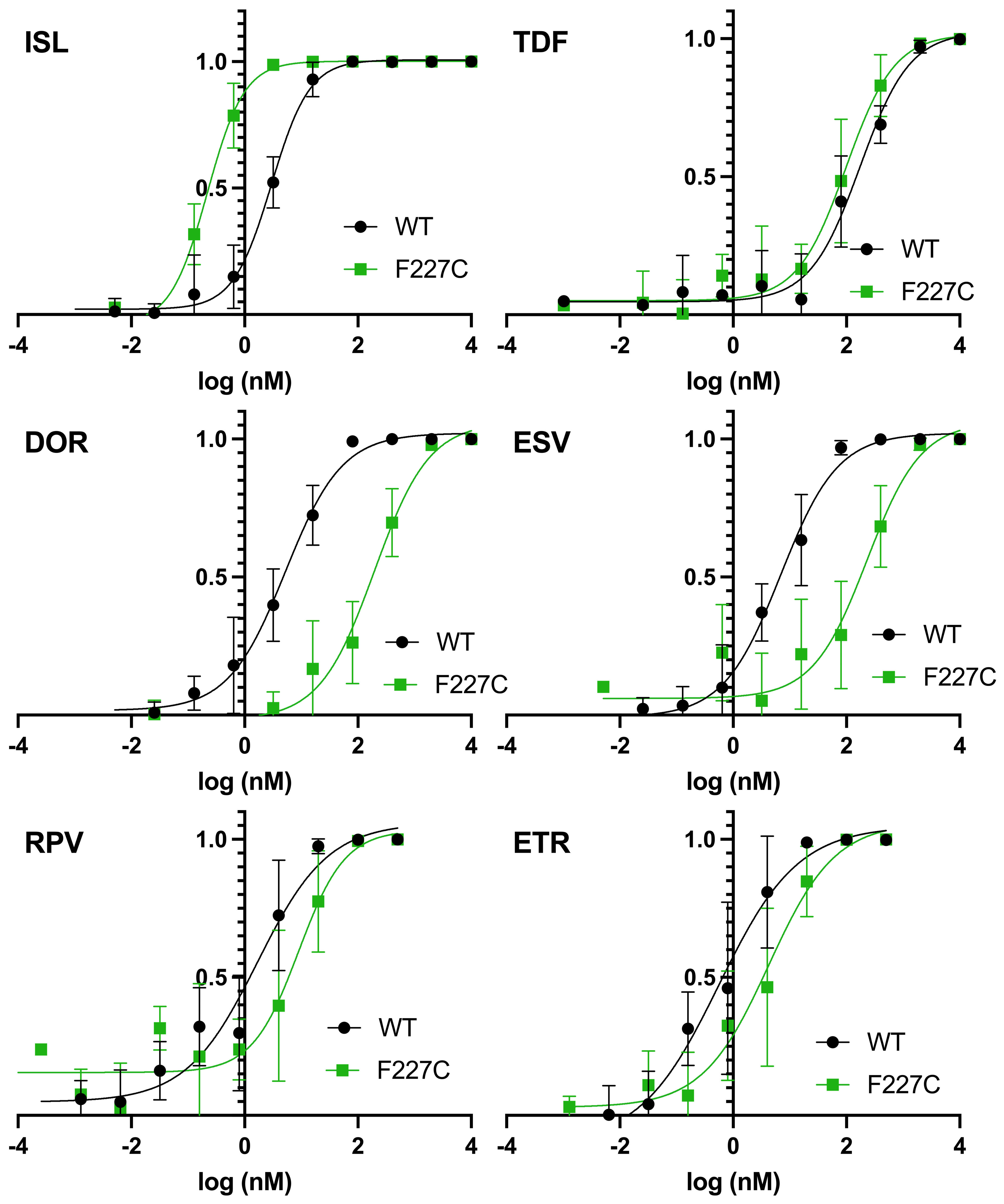

**Supplemental Figure 1. Dose-response curves of pNL4-3 ∆ env VSVG pseudotype WT and RT_F227C_ virus for RT Inhibitors: ISL, TDF, DOR, ESV, RPV, ETR.** TZM-GFP cells were pretreated with antivirals and infected after 24 h. GFP-positive cells (infected cells) were counted in various concentrations of inhibitor. Dose-response curves were produced for each mutant using Prism and the EC_50_s calculated and plotted (n=3-5).
