## Supplemental Figure 2 for "Structural and Mechanistic Basis of F227C-Mediated Hypersusceptibility to Islatravir in HIV-1 Reverse Transcriptase"

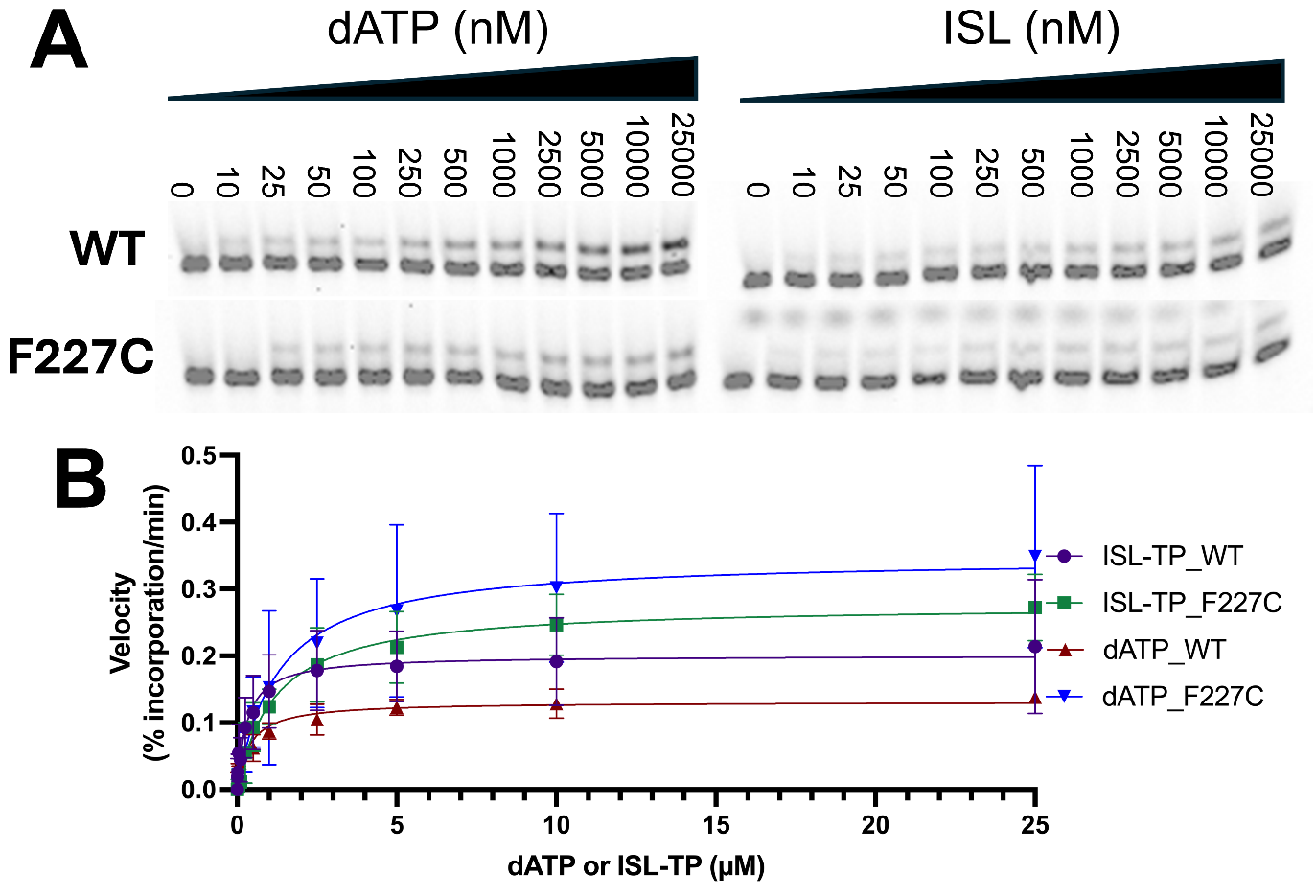

**Supplemental Figure 2. Incorporation kinetics of dATP and ISL-TP by WT and F227C HIV-1 RT.** (A) Representative primer extension gels showing incorporation of dATP or ISL-TP into the Td_31_/Pd_18_-P_0_ substrate (see Figure 2 for sequence) by RT_WT_ and RT_F227C_ across increasing substrate concentrations. Reactions were performed under steady-state conditions and resolved on a 15% polyacrylamide, 7 M urea denaturing gel. (B) Steady-state kinetic analysis of dATP and ISL-TP incorporation by WT and F227C RT. Incorporation velocity (µM/min) was determined from percent product formation over time and plotted as a function of substrate concentration. Data were fit to the Michaelis–Menten equation to determine steady-state kinetic parameters. Error bars represent standard deviation from at least three independent experiments.
