## Supplemental Figure 3 for "Structural and Mechanistic Basis of F227C-Mediated Hypersusceptibility to Islatravir in HIV-1 Reverse Transcriptase"

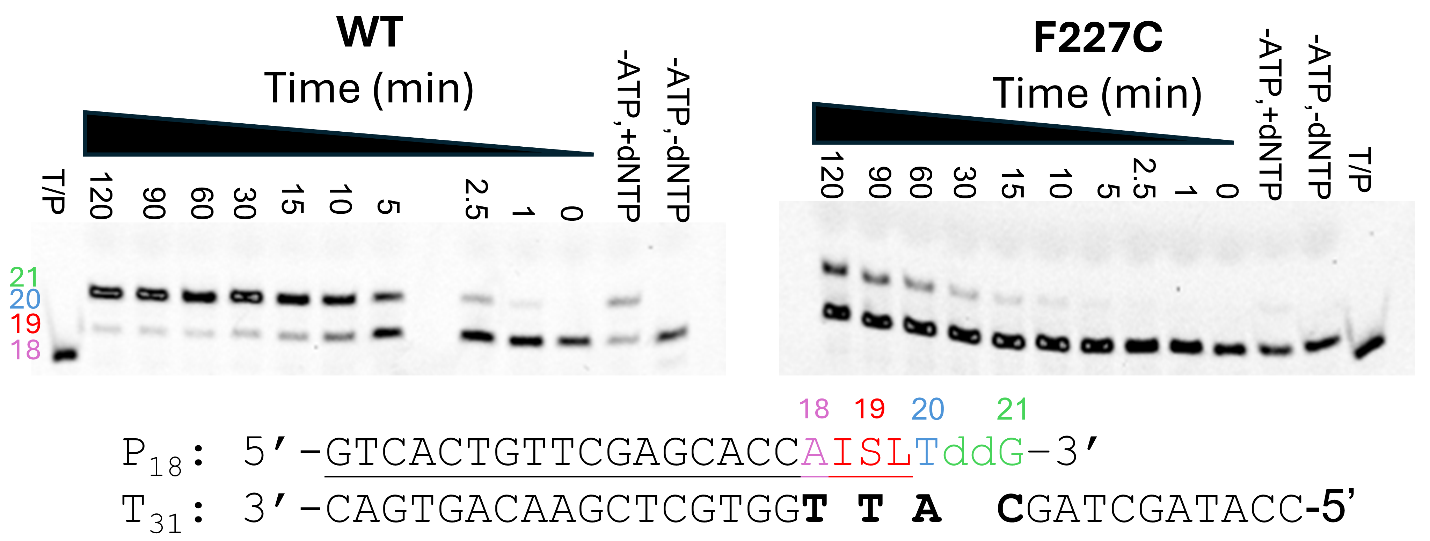

**Supplemental Figure 3. ATP-dependent rescue of ISL-MP-terminated primers by HIV-1 RT_WT_ and RT_F227C_. (**A) ATP-dependent rescue and extension of Td_31_/Pd_18_-P_0-ISL-MP_ by RT_WT_ and RT_F227C_. Purified Td_31_/Pd_18_-P_0-ISL-MP_ substrate was incubated with RT_WT_ or RT_F227C_ in the presence of 10 mM MgCl₂, 3.5 mM ATP, 100 µM dATP, 0.5 µM dTTP, and 10 µM ddGTP at 37°C. Aliquots were removed at the indicated time points, and reactions were quenched before analysis on a 15% polyacrylamide, 7 M urea denaturing gel. Representative gels from three independent experiments are shown.
