## Supplemental Figure 4 for "Structural and Mechanistic Basis of F227C-Mediated Hypersusceptibility to Islatravir in HIV-1 Reverse Transcriptase"

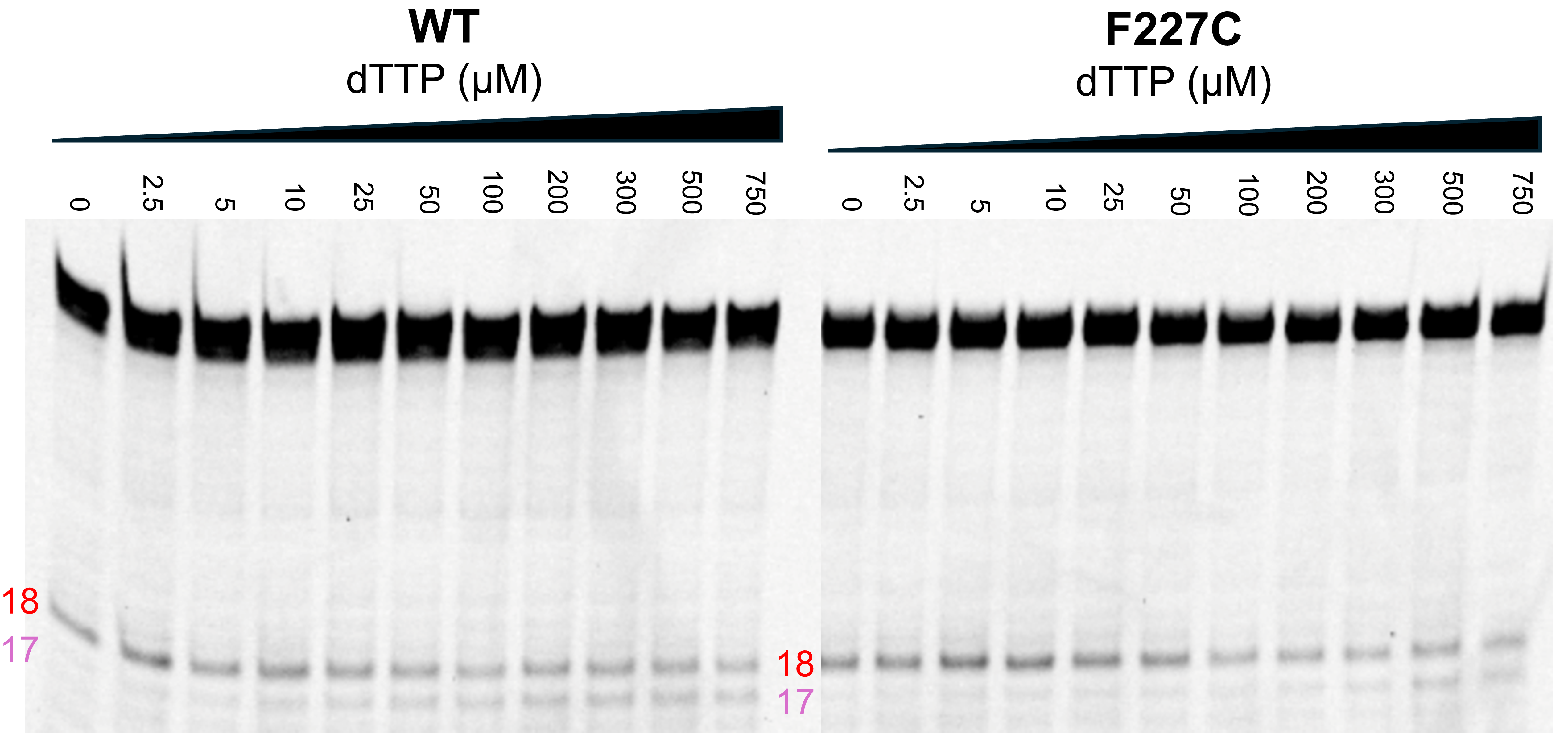

**Supplemental Figure 4. Effect of the F227C mutation on the translocation state of HIV-1 RT bound to T/P terminated with ISL-MP.** The translocation state of HIV-1 RT following ISL-MP incorporation was analyzed using site-specific Fe²⁺ footprinting. Td_43_/Pd_30-ISL-MP_ substrate (100 nM) containing a 5′-Cy3-labeled DNA template was incubated with RT_WT_ or RT_F227C_ (600 nM) in the presence of increasing concentrations of the next incoming nucleotide, dTTP. Complexes were treated with 1 mM ammonium iron sulfate for 5 min and resolved on a 7 M urea denaturing polyacrylamide gel. Cleavage at position −18 corresponds to the pre-translocation state, whereas cleavage at position −17 indicates the post-translocation state. Representative gels from three independent experiments are shown.
