## Supplemental Figure 5 for "Structural and Mechanistic Basis of F227C-Mediated Hypersusceptibility to Islatravir in HIV-1 Reverse Transcriptase"

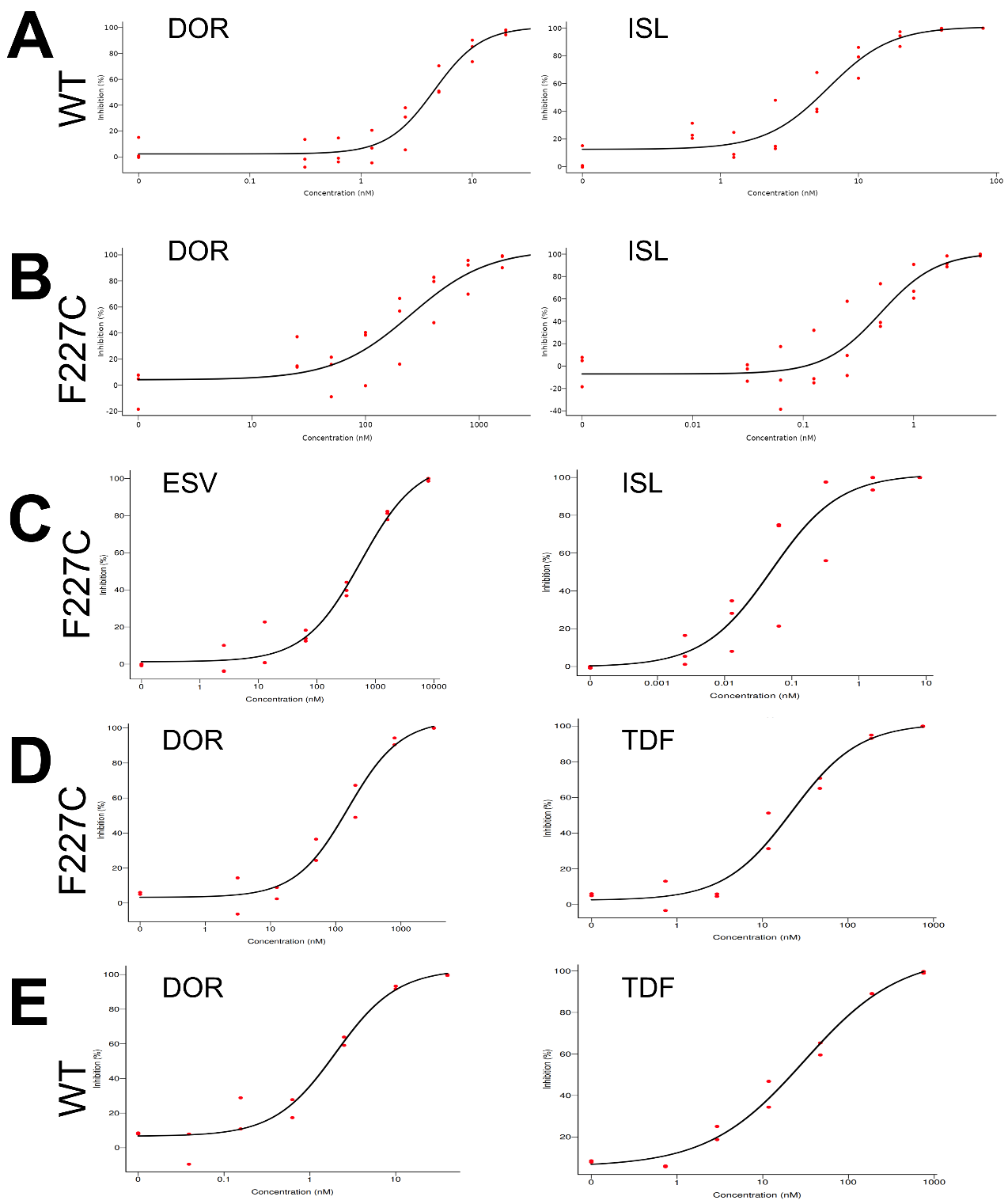

**Supplemental Figure 5. Representative dose-response curves from synergy experiments against WT and F227C HIV-1.**
Representative dose-response curves for (A,B) DOR + ISL, (C) ESV + ISL, and (D,E) DOR + TDF generated by SynergyFinder against WT and F227C HIV-1. TZM-GFP cells were preincubated with serially diluted inhibitors for 24 h before infection with NL4-3Δenv WT VSV-G pseudotyped virus in the presence of DEAE-dextran (1 µg/mL final concentration). Following an additional 48 h incubation, GFP-positive cells were quantified using a Cytation 5 imaging system and Gen5 v3.15 software. Percent inhibition values were normalized to no-drug controls. Data represent 2–4 independent experiments, each performed with technical duplicates.
