## Supplemental Figure 6 for "Structural and Mechanistic Basis of F227C-Mediated Hypersusceptibility to Islatravir in HIV-1 Reverse Transcriptase"

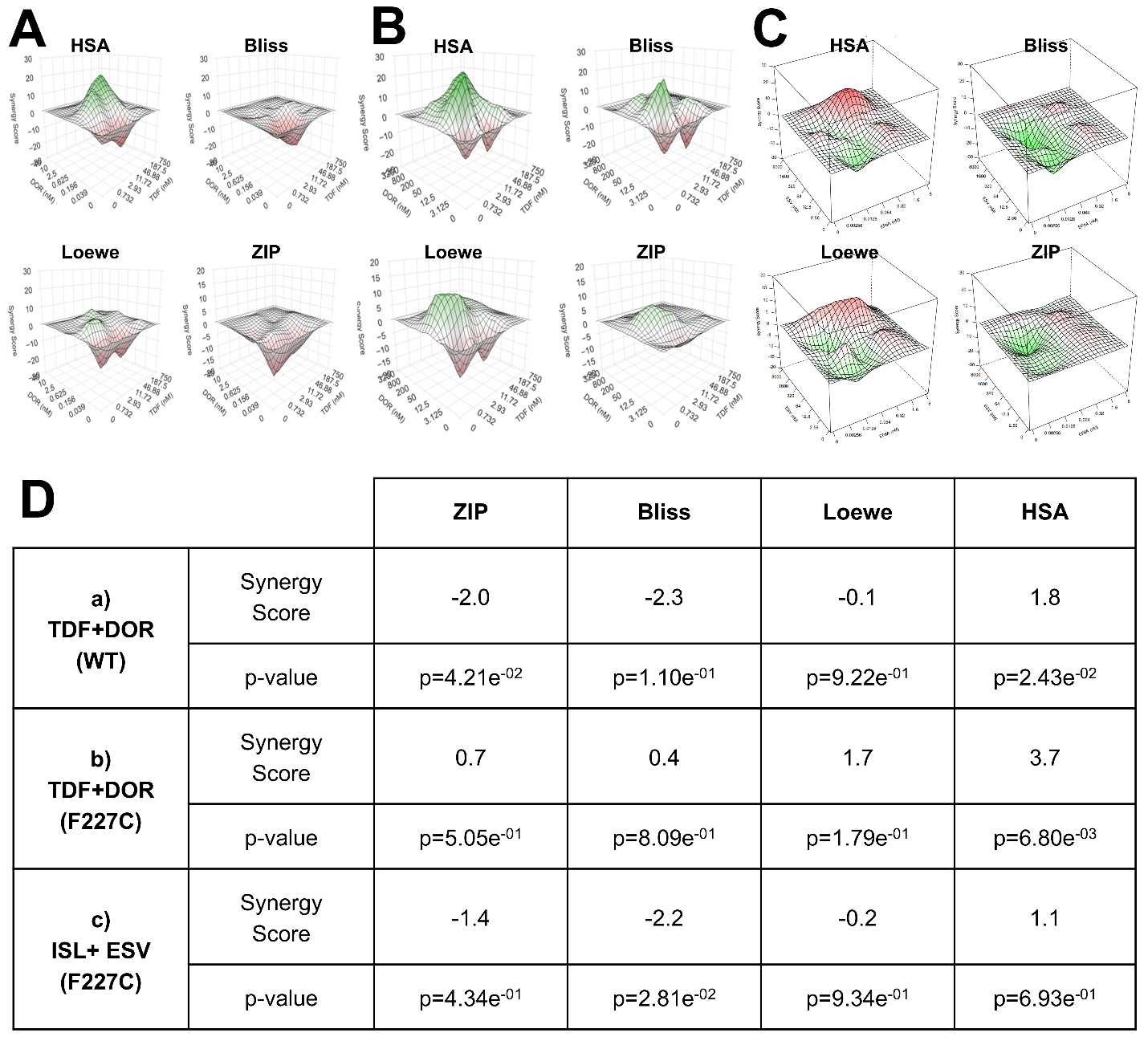

**Supplemental Figure 6. Combinations of NRTIs/NRTTIs and NNRTIs demonstrate predominantly additive antiviral interactions in WT and F227C HIV-1.** Three-dimensional synergy landscapes for combinations of NRTIs or NRTTIs with NNRTIs against WT and F227C HIV-1 NL4-3Δenv VSV-G pseudotyped virus were generated using the HSA, Bliss, Loewe, and ZIP reference models in SynergyFinder Plus. (A) TDF + DOR against the WT virus. (B) TDF + DOR against the F227C virus. (C) ISL + ESV against the F227C virus. Green regions indicate synergistic interactions, whereas red regions indicate antagonistic interactions. Synergy scores between −10 and 10 were interpreted as additive interactions, whereas scores >10 or <−10 indicate synergistic or antagonistic interactions, respectively. (D) Average synergy scores and associated P values for each reference model. Overall, all drug combinations exhibited predominantly additive interactions with localized regions of synergy in both WT and F227C viral backgrounds. Experiments were performed in duplicate or triplicate with averaged technical replicates.
