## Supplemental Table 1 for "Structural and Mechanistic Basis of F227C-Mediated Hypersusceptibility to Islatravir in HIV-1 Reverse Transcriptase"

**Supplemental Table 1. X-ray data collection and refinement statistics (PDB ID: XXXX)**

| **Data collection** | APS 22-ID, Dectris Eiger 16 M |
| --- | --- |
| Wavelength (Å) | 1.00 |
| Resolution (Å) | 1.8 (1.91-1.8)^a^ |
| Space group | *P*2_1_ |
| Cell dimensions |  |
| *a*, *b*, *c* (Å) | 83.99, 100.68, 92.49 |
| *β* (°) | 113.95 |
| Observed reflections | 891,349 |
| Unique reflections | 128,423 (20,517) |
| Redundancy | 6.9 |
| Completeness (%) | 99.0 (98.2) |
| R_meas_^b^ | 0.047 (1.25) |
| CC_1/2_ | 99.9 (85.2) |
| Avg I/σ | 18.2 (1.35) |
| **Refinement statistics** |  |
| Resolution (Å) | 43.3-1.8 |
| No. of reflections (working) | 128,091 |
| No. of reflections (test) | 6,295 |
| R_work_^c^ | 0.186 |
| R_free_^d^ | 0.214 |
| All-atom clashscore | 12.7 |
| Overall B value (Å^2^) | 58.12 |
| Wilson B value (Å^2^) | 39.7 |
| Ramachandran plot (%)^e^ |  |
| Favored | 96.64 |
| Allowed | 2.63 |
| Disallowed | 0.74 |
| RMSD Bond length (Å) | 0.007 |
| RMSD Angle (°) | 0.802 |
| ^a^ Values in parentheses are for the outer resolution shell. | |
| ^b^ R_meas_ = Σ*_hkl_* $\sqrt{\frac{n}{n-1}}$ $\sum_{j=1}^{n} \vert I_{hkl,j}- <I_{hkl}>\vert$ / Σ*_hkl_* Σ*_j_ I_hkl,j_*. | |
| ^c^ R_work_ = Σ*_hkl_* \|F*_obs_* – F*_calc_*\| / Σ*_hkl_* \|F_obs_\|. | |
| ^d^ R_free_ = R_work_, except 5% of the data excluded from the refinement. | |
| ^e^ Evaluated by MolProbity (Williams, C. J., *et al.* 2018. *Protein Sci* 27:293). | |
